# Differentiation drives widespread rewiring of the neural stem cell chaperone network

**DOI:** 10.1101/2020.03.05.976068

**Authors:** Willianne I. M. Vonk, T. Kelly Rainbolt, Patrick T. Dolan, Ashley E. Webb, Anne Brunet, Judith Frydman

## Abstract

Neural stem and progenitor cells (NSPCs) are critical for continued cellular replacement in the adult brain. Life-long maintenance of a functional NSPC pool necessitates stringent mechanisms to preserve a pristine proteome. We find that the NSPCs chaperone network robustly maintains misfolded protein solubility and stress resilience through high levels of the ATP-dependent chaperonin TRiC/CCT. Strikingly, NSPC differentiation rewires the cellular chaperone network, reducing TRiC/CCT levels and inducing those of the ATP-independent small heat shock proteins (sHSPs). This switches the proteostasis strategy in neural progeny cells to promote sequestration of misfolded proteins into protective inclusions. The chaperone network of NSPCs is more effective than that of differentiated cells, leading to improved management of proteotoxic stress and amyloidogenic proteins. However, NSPC proteostasis is impaired by brain aging. The less efficient chaperone network of differentiated neural progeny may contribute to their enhanced susceptibility to neurodegenerative diseases characterized by aberrant protein misfolding and aggregation.

## Introduction

Protein homeostasis (proteostasis) mechanisms in the brain are essential to prevent protein misfolding and aggregation-linked neurodegenerative diseases (Labbadia and Morimoto, 2015; Margulis and Finkbeiner, 2014; Morimoto and Cuervo, 2014; Taylor and Dillin, 2011). Adult brain plasticity and repair depends on neural stem and progenitor cells (NPSCs), which can both self-renew and differentiate into multiple neural cell lineages, thus facilitating learning, memory, cell replacement and repair after injury (Aimone et al., 2006; Arvidsson et al., 2002; Bonaguidi et al., 2011; Dupret et al., 2008; Spalding et al., 2013). Brain aging dramatically impairs the regenerative potential of the NSPC pool (Apple et al., 2017; Luo et al., 2006; Maslov et al., 2004; Rando, 2006; Vilchez et al., 2014). The decreased ability of aged NSPCs to replace damaged or lost neurons may contribute to age-dependent neurodegenerative diseases, such as Alzheimer’s, Parkinson’s, and Huntington’s disease (HD), and amyotrophic lateral sclerosis (ALS). Additionally, aging itself leads to reduced proteostasis capacity across tissues, including the brain, which further enhances formation of potentially neurotoxic, misfolded proteins (Brehme et al., 2014; Hipp et al., 2019; Morimoto and Cuervo, 2014; Taylor and Dillin, 2011).

Life-long maintenance of a functional NSPC pool necessitates stringent mechanisms that promote genome and proteome integrity (Burkhalter et al., 2015; Llorens-Bobadilla et al., 2015; Shin et al., 2015). The complex proteostasis network (PN) maintaining the proteome consists of different molecular chaperones and clearance pathways. These contribute distinctly to cellular proteostasis (Freilich et al., 2018) but together determine polypeptide fate throughout the life of a protein (Jayaraj et al., 2019; Sala et al., 2017). How the distinct PN components recognize a misfolded protein and decide whether to refold, degrade or sequester it into an inclusion is poorly understood; particularly in stem cells and other highly specialized cell types. Previous studies showed that proteasome levels are high in stem cells, while autophagy is induced upon differentiation (Aron et al., 2018; Ho et al., 2017; Kenyon, 2010; Koyuncu et al., 2018; Tang and Rando, 2014; Vilchez et al., 2012; Vilchez et al., 2014). The function of these changes in the context of overall proteostasis and proteome maintenance remains poorly understood. Here, we compare the proteostasis networks and strategies of adult NSPC with their differentiated progeny. We find that differentiation induces widespread remodeling of the chaperone networks of NSPCs, fundamentally changing their strategies of proteome maintenance. Differentiation dramatically downregulates the ATP-dependent chaperonin TRiC/CCT, and induces the ATP-independent small heat shock proteins (sHSPs). While high TRiC in NSPCs robustly maintains protein solubility and suppresses amyloid aggregation, sHSPs induced in differentiated progeny promote a switch in proteostasis strategy to rely on chaperone-dependent sequestration of misfolded proteins into inclusions. Notably, the robust NSPC chaperone network provides effective stress resilience and cellular fitness, while differentiated cells are more vulnerable to stress and amyloid deposition. Our findings reveal an unexpected complex tuning of proteostasis maintenance, whereby the chaperone composition of a cell determines its preferred protein quality control (PQC) strategy.

## Results

### Distinct proteostasis capacities of NSPCs and their differentiated progeny

To identify proteostasis mechanisms maintaining cellular fitness in adult NSPCs, we isolated Sox2^+^/Nestin^+^ NSPCs from the two stem cell niches in adult mouse brain, the subgranular zone (SGZ) of the hippocampal dentate gyrus and the subventricular zone (SVZ) of the anterior lateral ventricle. In culture, NSPCs are proliferative and retain their ability to both self-renew or differentiate into a mixed population of three main neural lineages, i.e. astrocytes, neurons, and oligodendrocytes, representative of the mixed *in vivo* differentiated cell population of the adult brain (Figures 1A; S1A, B) (Artegiani et al., 2017; Bonaguidi et al., 2011; Walker and Kempermann, 2014). Comparing the ability of the activated, proliferative NSPCs and their differentiated progeny to withstand various proteotoxic stresses revealed dramatic differences in their cellular resistance to both oxidative stress and heat shock. While adult NSPCs were highly resistant to proteotoxic stress, the mixed population of differentiated neural cell lineages was uniformly very sensitive to these treatments (Figures 1B, S1C).

**Figure 1.**
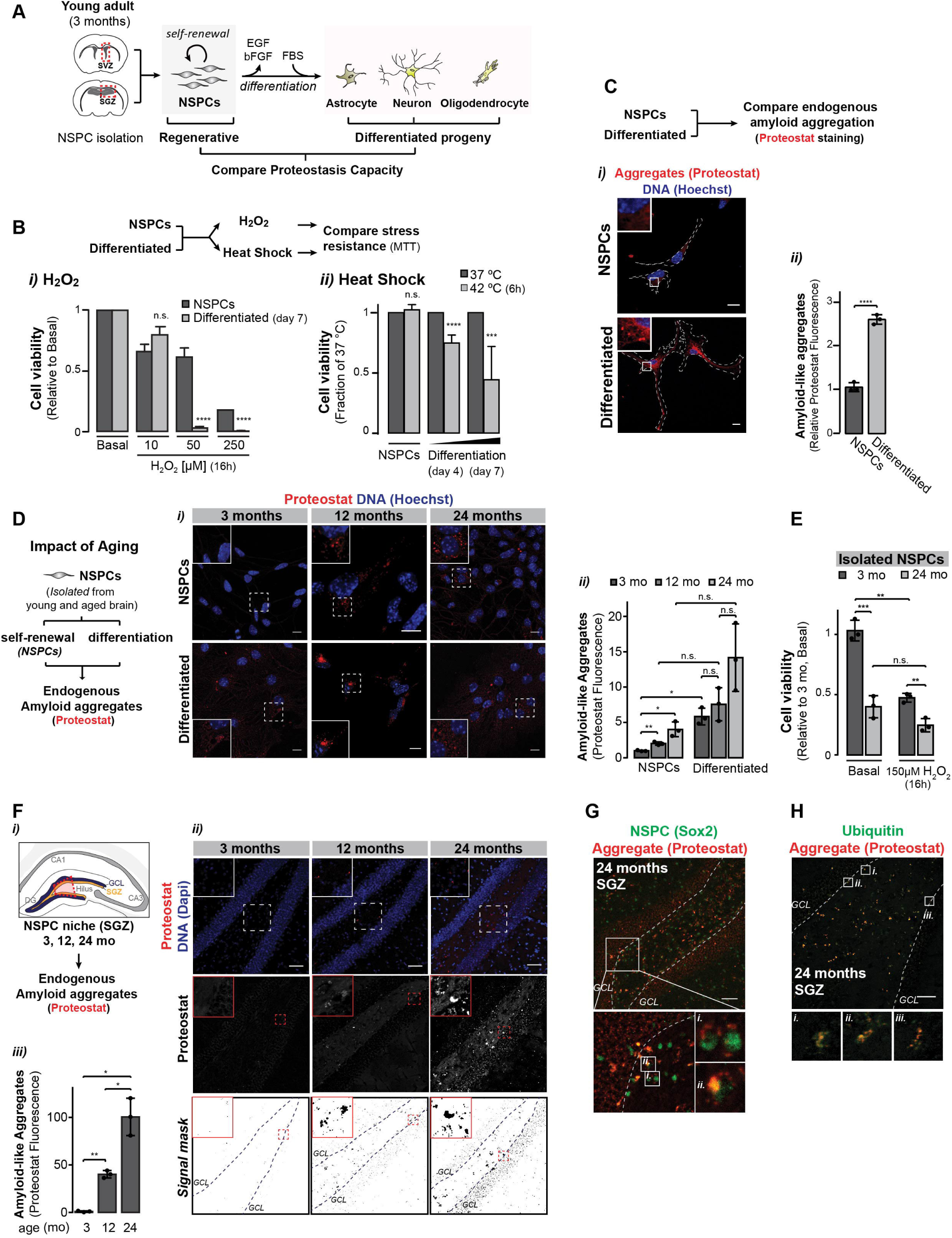
Robust proteostasis ability of NSPCs *vs*. differentiated lineages. **A**, Regenerative NSPCs, isolated from the SVZ and SGZ stem cell niches of the young adult (3 month old) mouse brain, were differentiated into three main neural lineages (i.e. astrocytes, neurons, oligodendrocytes). **B**, Stress resilience of NSPCs and their differentiated progeny, determined upon exposure to (***i***) oxidative stress ([H_2_O_2_]) and (***ii***) heat stress. Viability was normalized for basal conditions. Basal unstressed conditions yield lower MTT reduction into formazan by differentiated progeny relative to NSPCs (metabolic activity of differentiated cells was attenuated by 49 ± 3% compared to NSPCs (Figure 5E)). n.s., non-significant, *** p < 0.001, **** p < 0.0001. **C**, (***i***) Proteostat fluorescence and nuclear DNA (Hoechst). Scale bars: 10 µm; traces: cell outlines. Insets: Magnification of Proteostat^+^ deposits. (***ii***) Quantified Proteostat fluorescence (mean ± SD). **** p < 0.0001. **D**, (***i***) Proteostat and Hoechst in NSPCs isolated from brains of 3, 12, and 24 month old mice and differentiated progeny cells. Insets: Magnification of Proteostat^+^ deposits. (***ii***) Total Proteostat fluorescence (mean ± SD; quantified by fluorimeter). Proteostat fluorescence of young NSPCs is set at 1. * p < 0.05; ** p < 0.01; n.s., non-significant. **E**, Relative cell viability (mean ± SD) of young and old NSPCs under basal and oxidative stress conditions. Viability for young NSPCs at basal conditions is set at 1. ** p < 0.01; *** p < 0.005; n.s. non-significant. **F**, (***i***) Schematic overview of SGZ region in the dentate gyrus (DG) of the hippocampus. GCL: granular cell layer. (***ii***) Proteostat^+^ deposits in the SGZ (red marked area in ***i***) of 3, 12, and 24 month old mouse brains. Bottom row: Proteostat fluorescence signal masks. Scale bars: 50 µm. (***iii***) Relative Proteostat fluorescence in hippocampal brain sections (whole frame; mean ± SD). Fluorescence at 3 months is set to 1. * p < 0.05, ** p < 0.01. Insets: Magnification of Proteostat^+^ depositions. See Figure S1D for additional ages. **G, H**, Sox2^+^ (**G**) or ubiquitin^+^ (**H**)/Proteostat^+^ inclusions in the DG of the aged brain. Scale bar: 50 µm. Insets: Magnification of Proteostat^+^ deposits.

Impaired proteostasis is linked to formation of amyloid aggregates, typical of aging-linked neurodegenerative diseases (Hipp et al., 2019; Morimoto and Cuervo, 2014; Taylor and Dillin, 2011). We examined if unstressed NSPCs and their differentiated progeny could equally prevent amyloid aggregation of endogenous proteins using Proteostat, a dye that fluoresces upon selectively binding to amyloid-like protein deposits (Shen et al., 2011). NSPCs were almost completely devoid of aggregates; in contrast, Proteostat^+^ inclusions were abundant in unstressed differentiated cells (Figure 1C). We conclude that proteostasis capacity is much higher in NSPC than in differentiated cells, endowing NSPCs with robust stress resistance and preventing amyloid aggregation.

We next examined the impact of aging on NSPC proteostasis and fitness. Aging is associated with a decline in proteostasis capacity (Josefson et al., 2017; Leeman et al., 2018; Morimoto and Cuervo, 2014; Schmidt and Finley, 2014; Taylor and Dillin, 2011; Vilchez et al., 2014) as well as depletion of the functional stem cell pool (Apple et al., 2017; Artegiani et al., 2017; Kalamakis et al., 2019; Luo et al., 2006; Maslov et al., 2004; Vilchez et al., 2014). We examined the presence of Proteostat^+^ deposits in NSPCs isolated from either young adult, middle-aged, or aged mouse brains (i.e. 3, 12, and 24 months, respectively; hereafter referred to as young, middle-aged, or aged NSPCs). Young NSPCs were almost completely devoid from Proteostat^+^ puncta, but their abundance was progressively enhanced during brain aging (Figure 1D). This age-dependent decline in NSPC proteostasis capacity correlated with reduced cellular fitness, which was exacerbated upon exposure to proteotoxic stress (Figure 1E). To account for possible effects of NSPC isolation and culture on cellular proteostasis, we directly examined the presence of Proteostat^+^ deposits in mouse SGZ brain sections. While SGZ stem cells in the brains of young animals were devoid of amyloid aggregates, there was an age-dependent increase in amyloid aggregates in SGZ stem cells of middle-aged and aged mice (Figures 1F, S1D). These Proteostat^+^ deposits co-localized with Sox2^+^ stem cells (Figure 1G) and were ubiquitin-positive (Figures 1H, S1E). Of note, Proteostat^+^/ubiquitin^+^ protein inclusions characterize misfolding-linked neurodegenerative pathologies (Shen et al., 2011). These studies indicate that NSPCs have high proteostasis capacity which declines during aging, leading to increased amyloid aggregation.

### Differentiation alters NSPC management of polyQ-expanded Huntingtin

Next, we asked whether the distinct proteostasis capacities of adult NSPCs and their progeny also affect their management of HD-causing, polyQ-expanded Huntingtin (Htt; (Escusa-Toret et al., 2013; Kaganovich et al., 2008; Polling et al., 2014)). To this end, we analyzed the subcellular distribution of HttQ97 exon1-GFP in both NSPCs and their differentiated progeny. In all three differentiated neural lineages, HttQ97-GFP exhibited a punctate distribution; consistent with sequestration into inclusions (Figures 2A, B). Strikingly, despite its high aggregation propensity, HttQ97-GFP was mainly diffusely localized in NSPCs, with only very few inclusions (Figure 2A). To examine if this altered HttQ97-GFP localization is linked to the enhanced NSPC proteostasis, we next compared the cellular toxicity of HttQ97-GFP in NSPCs and differentiated cells. Exogenous expression of pathogenic Htt significantly increased cell death in the differentiated population, but did not affect NSPC viability (Figure 2C). These findings suggest that NSPCs are more resilient than differentiated cells to HttQ97-GFP, while differentiated cells are less effective at managing polyQ misfolding and toxicity.

**Figure 2.**
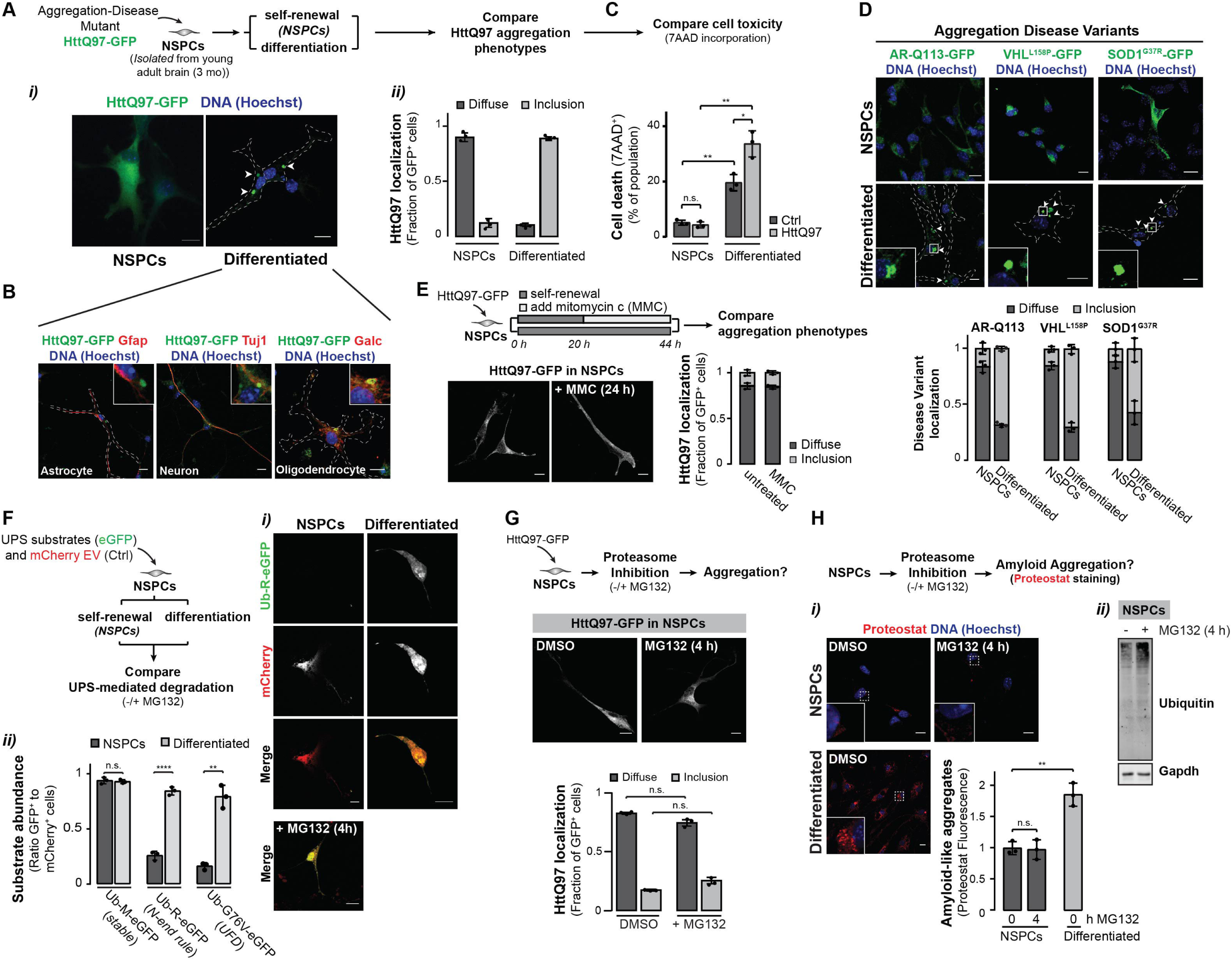
High capacity of NSPC to maintain proteome integrity. **A**, (***i***) HttQ97-GFP, transfected into NSPCs and maintained under self-renewal or differentiation conditions, and DNA (Hoechst). Traces: cell outlines; arrows: HttQ97 aggregates. (***ii***) Quantification (mean ± SD) of cells with diffusely *vs*. aggregated HttQ97-GFP (≥ 150 cells/condition/experiment). **B**, HttQ97-GFP, DNA (Hoechst), and specific markers for differentiated lineages. Traces: cell outlines. Insets: Magnification of perinuclear aggregation. **C**, Dead (7AAD^+^) cells (mean ± SD) in NSPC and differentiated HttQ97-GFP^−^ (Ctrl) and HttQ97-GFP^+^ cell populations. n.s., non-significant, * p < 0.05, ** p < 0.01. **D**, AR-Q113, VHL^L158P^, and SOD1^G37R^-GFP fusion proteins and DNA (Hoechst) in NSPCs and differentiated progeny. Traces: cell outlines; arrows: aggregates. Insets: Magnification of inclusion. Quantification (mean ± SD) of diffusely localized *vs*. aggregated proteins (≥ 100 cells/condition/experiment). **E**, HttQ97-GFP in control and MMC-treated (1µg/ml, 24 h) NSPCs. Scale bars: 10 μm. Quantification (mean ± SD) of diffuse *vs*. aggregated HttQ97-GFP (≥ 60 cells/condition/experiment). See also Figure S2E. **F**, (***i***) Ub-R-eGFP and mCherry-empty vector control plasmid (transfection control) in NSPCs and differentiated progeny. UPS-mediated clearance was determined at basal conditions and upon proteasomal blockage (+ MG132; 15 µM, 4 h) by relative quantification of GFP^+^/mCherry^+^ cells. (***ii***) Graph (mean ± SD) quantifies Ub-R-eGFP as well as Ub-G76V-eGFP and their stable counterpart Ub-M-eGFP (≥ 100 cells/condition). ** p < 0.01; **** p < 0.0001; n.s., non-significant. **G**, HttQ97-GFP in NSPCs at vehicle control-treated (DMSO) or proteasome-inhibited (MG132; 15 µM, 4 h prior to fixation) conditions. Graph (mean ± SD) quantifies diffusely localized *vs*. aggregated HttQ97-GFP (≥ 60 transfected cells/condition/experiment). n.s., non-significant. **H**, *i)* Amyloid-like aggregates (Proteostat^+^) and DNA (Hoechst) in NSPCs and differentiated progeny at control (DMSO) or proteasome-inhibited conditions (MG132; 15 µM, 4 h prior to fixation). Insets: Magnification of Proteostat^+^ deposits. Quantification (mean ± SD) shows Proteostat fluorescence. ** p < 0.01; n.s. non-significant. See also Figure S2H. *ii*) Immunoblot for Ubiquitin in NSPCs at basal or proteasome-inhibited conditions (MG132; 15 µM, 4 h prior to fixation). Gapdh serves as loading control.

To extend the comparison of the proteostasis capacities of NSPCs *vs*. the mixed population of differentiated lineages, we examined the localization of a panel of disease-linked, aggregation-prone mutants with distinct properties and chaperone requirements: the polyQ-expanded Androgen Receptor (AR-Q113), the cancer-associated Von Hippel-Lindau tumor suppressor (VHL^L158P^), and the ALS-linked superoxide dismutase 1 (SOD1^G37R^) (Table S1; (Escusa-Toret et al., 2013; Sontag et al., 2014)). In all cases, similar to HttQ97-GFP, the misfolded protein was sequestered into inclusions in differentiated cells but remained predominantly diffusely localized in NSPCs (Figure 2D). We conclude that the NPSC proteostasis circuits maintain solubility of a wide range of disease-linked, misfolded proteins while promoting enhanced resistance to distinct proteotoxic stresses. In contrast, differentiated lineages tend to sequester misfolded proteins into inclusions and are more sensitive to proteotoxic stress.

We next examined if NSPCs rely on their high division rates to reduce aggregate abundance and protect cells from stress by diluting misfolded proteins. Since aggregation can be concentration dependent, we first compared whether HttQ97-GFP transfection prior to or post-differentiation altered the expression or distribution of the misfolded protein. HttQ97 localization and inclusion abundance were similar for both populations of differentiated cells (Figure S2A). HttQ97-GFP expression in the differentiated progeny was comparable to that in NSPCs, despite their distinct localization (Figure S2B). In an alternative approach, NSPC proliferation was blocked using mitomycin C (MMC), which should prevent dilution of damaged proteins. MMC treatment reduced the NSPC pool size without affecting its fitness (Figures S2C*ii*, D*ii*). Importantly, MMC treatment did not lead to the appearance of HttQ97-GFP inclusions nor increased the incidence of endogenous Proteostat^+^ aggregates (Figures 2E; S2C-E). Similar results were obtained upon prolonged incubation or increased concentrations of MMC (Figures S2C-E). We thus conclude that NSPCs maintain proteome solubility through their robust proteostasis capacity rather than through misfolded protein dilution during cell division.

Next, we examined the regulation of clearance pathways. Stem cells have been shown to exhibit high proteasome levels (Jang et al., 2014; Vilchez et al., 2012; Zhao et al., 2016). We examined whether efficient proteasomal clearance could explain the robust NSPC proteostasis. First, we assessed the ubiquitin-proteasome system (UPS) in NSPCs and their progeny. Clearance assessment of two distinct short-lived, GFP-fused UPS reporters for N-end rule or ubiquitin-fusion degradation (Dantuma et al., 2000) showed UPS substrates are efficiently degraded in NSPCs but accumulate in differentiated cells; indicative of low UPS activity (Figure 2F). Consistent with UPS-mediated clearance, treatment with proteasome inhibitor MG132 stabilized the unstable UPS substrates in NSPCs (Figure 2F*i*). Biochemical analyses further indicated that NSPCs contain much higher proteasome activity as well as 19S and 20S proteasomal complex expression than their progeny (Figures S2F, G). However, proteasome inhibition by MG132 in NSPCs did not lead to increased endogenous amyloid formation nor to formation of HttQ97-GFP puncta (Figures 2G, H; S2H). These experiments indicate that proteasome-mediated clearance cannot explain the solubility of the NSPC proteome.

### Transcriptomic analyses reveal rewiring of the NSPC chaperone network upon differentiation

To gain insights into how differentiation affects NSPC proteostasis, we used RNA sequencing (RNA-Seq) to monitor changes in the NSPC transcriptome following 7 days of differentiation (Figure 3A*i*, Table S2). As expected, differentiation reduced stem cell specific genes and enriched genes expressed in neural lineages (Figure 3A*ii*). Analysis of biological processes identified various pathways and categories enriched in NSPCs *vs*. their differentiated progeny (Table S3, Figure S3A). We focused on those contributing to proteostasis.

Cellular proteostasis relies on the interplay of protein synthesis with diverse chaperone classes and clearance systems (Figure 3B). Unlike what is observed upon differentiation of embryonic stem cells (ESCs), which leads to higher translation (Ingolia et al., 2011; Sampath et al., 2008; You et al., 2015), NSPC differentiation led to an overall decline in transcripts encoding ribosomal subunits, indicative of translation attenuation (Figures 3B, S3A). Direct measurements of bulk protein synthesis using fluorescent O-propargyl-puromycin (OPP) labeling of newly translated proteins confirmed translation is reduced in the neural progeny (Figures 3C, S3B). Paradoxically, higher translation rates enhance the load on the chaperone machinery and contribute to proteostasis dysfunction (Hansen et al., 2007; Sherman and Qian, 2013). Thus, the high NSPC translation rate cannot account for their high proteostasis capacity and stress resilience; if anything, it should present a higher hurdle to proteostasis maintenance.

**Figure 3.**
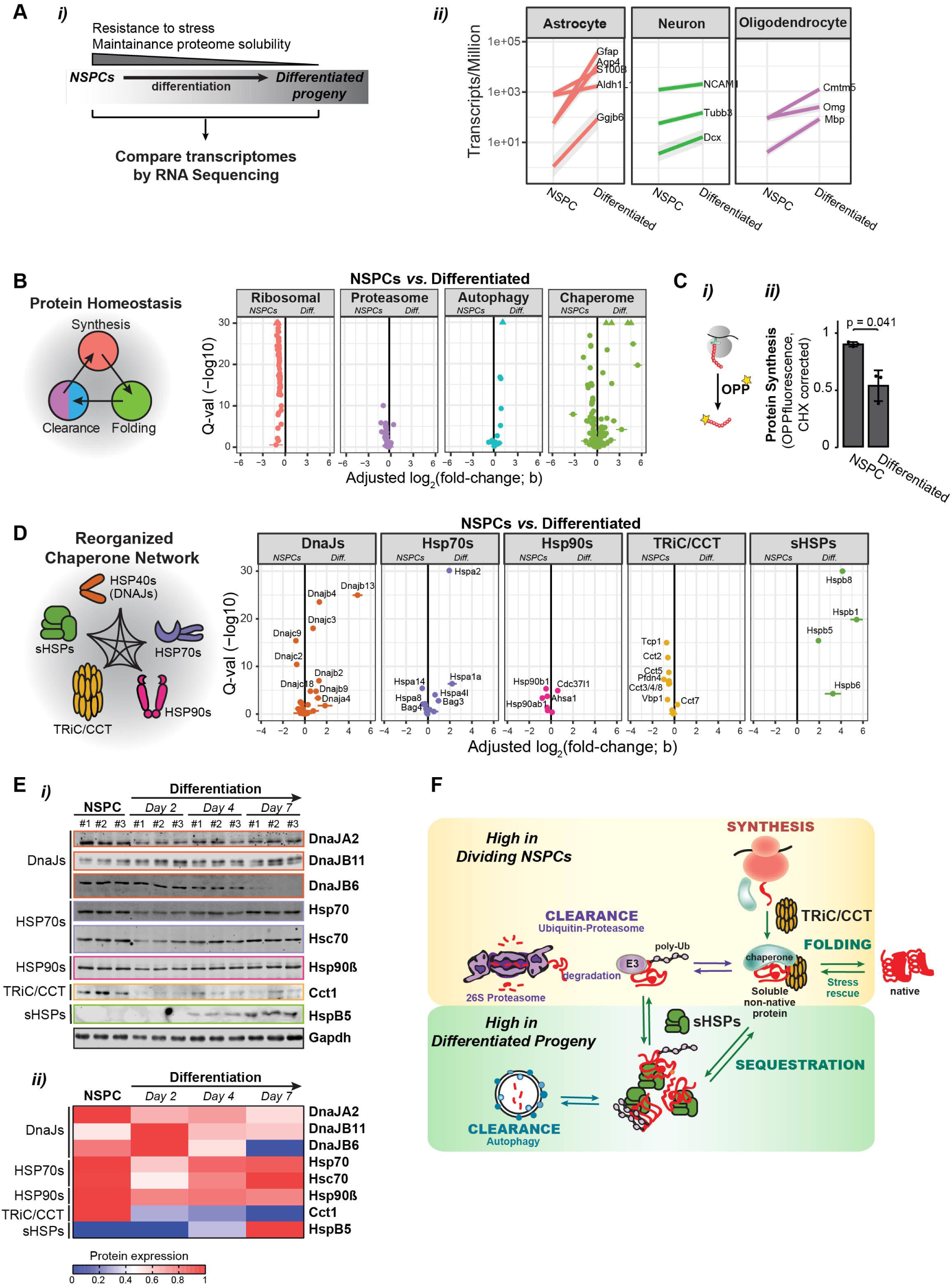
NSPC differentiation rewires cellular chaperone circuits. **A**, (***i***) The robust NSPC proteostasis capacity (i.e. high stress resistance, ability to maintain proteome solubility) is drastically impaired upon differentiation. The differential transcriptomes of NSPCs and their progeny was compared by RNA-Seq. (***ii***) Proof of concept: marked enrichment of transcripts specifically expressed in the differentiated lineages following NSPC differentiation. **B**, Cellular proteostasis relies on the balance of protein synthesis, folding, and clearance, mediated by molecular chaperones and degradation pathways acting during protein maturation. Volcano plots show differentially expressed ribosomal, proteasomal, autophagy or chaperome genes in differentiated progeny compared to NSPCs. X-axis: log2 (fold change); y-axis: –log10 of q-value. Triangles indicate q-values below 10^−30^. **C**, (***i***) Bulk rates of protein synthesis, determined by O-propargyl-puromycin (OPP) incorporation into newly translated proteins. (***ii***) Quantification (mean ± SD) of the relative protein synthesis rates in NSPCs and differentiated progeny. Value of NSPCs was arbitrary set at 1. p = 0.041. See Figure S3B for FACS plots. **D**, Volcano plots of differentially expressed genes of all five major cytosolic chaperone classes in differentiated cells compared to NSPCs. X-axis: log2 (fold change); y-axis: –log10 of q-value. **E**, (***i***) Immunoblots for indicated chaperones for three independent samples of NSPCs and early, intermediate, and fully differentiated cells. Gapdh serves as loading control; (***ii***) Heat map of immunoblot quantifications. **F**, Eukaryotic cells contain different chaperone classes guiding folding of non-native polypeptides or effecting quality control through protein refolding, degradation, and sequestration. We postulate a model in which high TRiC in NSPCs ensures maintenance of a soluble proteome, by TRiC binding to nascent and non-native polypeptides to promote their folding into native proteins or clearance by proteasome or autophagy pathways. NSPC differentiation remodels the cellular chaperone networks and induces sHSP expression. This may change the cellular proteostasis strategy and promote stabilization of non-native proteins by spatial sequestration into defined protective inclusions.

Next, we examined changes in clearance pathways. Consistent with previous data and our experimental observations (Figures 2F-H, S2F-H), multiple 19S and 20S proteasome subunit transcripts were enriched in NSPCs (Figures 3B, S3C). Analysis of differentially regulated ubiquitin pathway genes identified important UPS-requiring processes in NSPCs, including DNA replication and repair, in addition to stress resistance (Figures S3D-F). Differentiation led to upregulation of autophagy pathway genes (Figures 3B, S3G; and (Fimia et al., 2007; Vazquez et al., 2012)), suggesting a switch in clearance strategies. Of note, the experimentally measured autophagic flux in differentiated cells was not significantly increased compared to NSPCs (Figure S3G); this could arise from a substrate overload of autophagy due to the much lower proteasome activity levels.

We next analyzed the differential regulation of chaperone networks. RNA-Seq indicated a clear remodeling of the cellular chaperome upon NSPC differentiation (Figure 3B). We examined the differential expression of the five major cytoplasmic chaperone families with distinct functions, namely HSP40/DnaJ HSP70 co-factors, HSP70s, HSP90s, TRiC/CCT, and small HSPs (sHSPs) (Hartl et al., 2011; Powers and Balch, 2013; Yerbury et al., 2016) (Figure 3D). There were a few changes in expression of most Hsp70 and Hsp90 isoforms. Several specific Hsp70-cofactors of the DnaJ/Hsp40 and Bag families were either upregulated, down-regulated, or unchanged upon differentiation (Figure 3D). Most notably, NPSC differentiation distinctly affected two chaperone classes, TRiC/CCT and the sHSPs. For both, the RNA-Seq results were validated by RT-qPCR. All subunits from the ATP-dependent TRiC chaperonin, CCT1-CCT8 (Leitner et al., 2012), were downregulated in differentiated cells (Figures 3D, S4A), while several ATP-independent sHSPs were induced (Figures 3D, S4D). A *Cct1* promoter reporter further corroborated the transcriptional downregulation of *Cct* genes upon NSPC differentiation (Figure S4C). NSPC passage number did not affect the expression pattern of proteostasis genes, indicating that these changes are not influenced by *in vitro* culture (Figure S4B).

The alterations in chaperone expression during NSPC differentiation were next studied at the protein level using immunoblot analyses. We did not observe significant changes for the major HSP90 and HSP70 isoforms as well as HSP40/DnaJ isoforms DnaJA2 and DNAJB11 (Figure 3E). Of note, DnaJB6, which is unchanged at the RNA level, did decline at the protein level late during NSPC differentiation (Figure 3E). We also observed that TRiC subunits were abundantly expressed in NSPCs but declined dramatically during differentiation (e.g. Cct1 in Figure 3E). Conversely, several sHSPs were undetectable in NSPCs but strongly induced during differentiation (e.g. HspB5 in Figure 3E).

These data indicate that NSPC differentiation does not simply downregulate chaperones and clearance factors, but instead rewires the PN by differential expression of chaperone machineries with distinct modes of action. In particular, the ATP-dependent chaperonin TRiC is known to promote folding and suppress aggregation of newly translated proteins (Gestaut et al., 2019), while the ATP-independent sHSPs can stabilize misfolded proteins through sequestration in inclusions (Mogk et al., 2019). We thus hypothesized that differentiation-linked changes in the levels of these chaperone systems, likely in concert with additional chaperone co-factors and clearance pathways, underlie the distinct strategies by which NSPCs and differentiated cells manage misfolded proteins (Figure 3F).

### Differential expression of TRiC and sHSPs in NSPCs and differentiated progeny

To test our hypothesis, we examined further the changes in TRiC and sHSP expression upon NSPC differentiation. Immunoblot analyses confirmed that all eight Cct subunits were highly expressed in NSPCs, and dramatically reduced in differentiated cells (Figure 4A). Conversely, HspB1, HspB5, and HspB8 proteins were virtually undetectable in NSPCs but highly induced upon differentiation (Figures 4B, S4E). Other sHSPs (e.g. HspB3, HspB6) remained unaffected during differentiation (Figure 4B).

**Figure 4.**
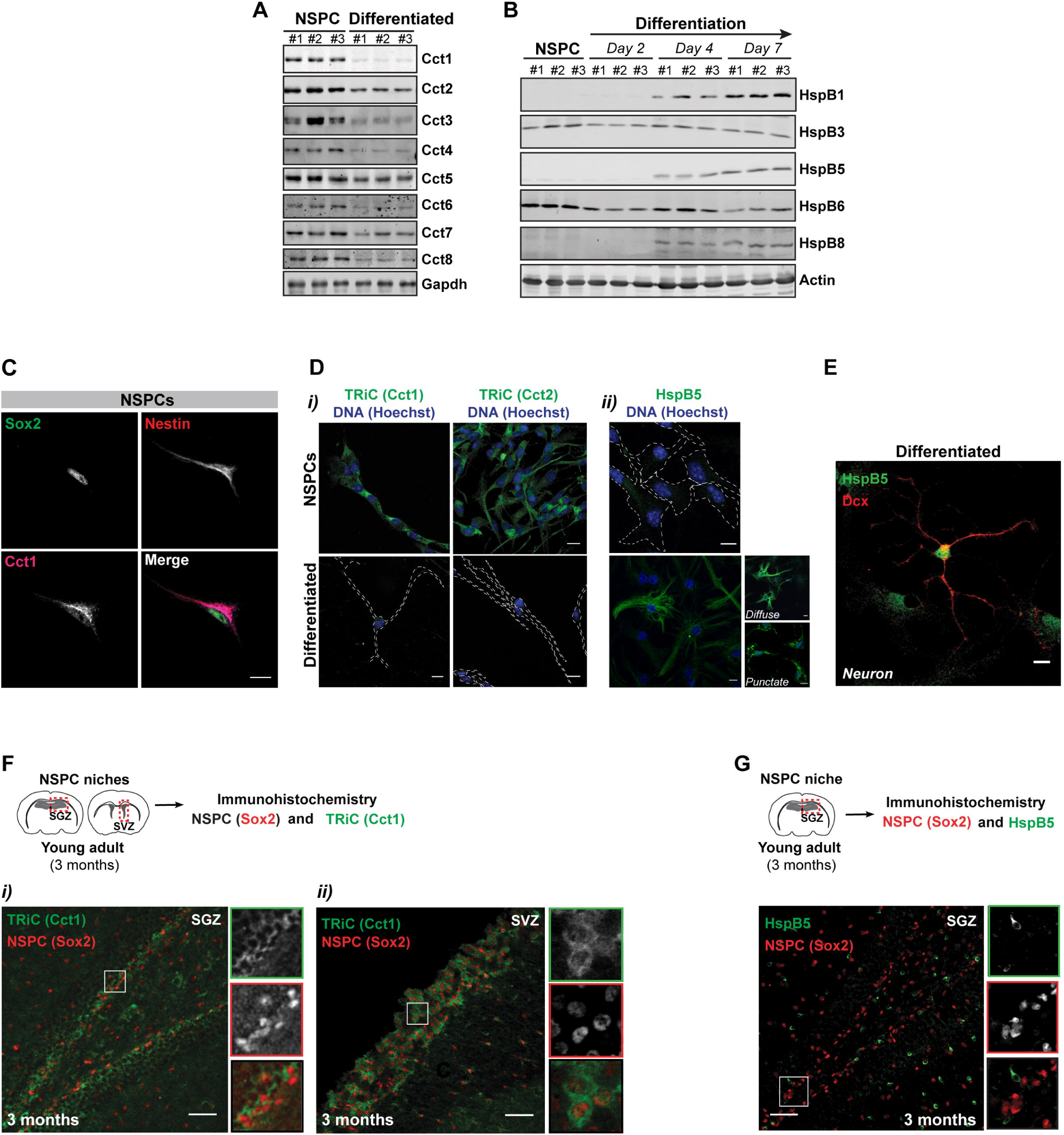
Differential expression of TRiC and HspB5 in NSPCs and differentiated progeny. **A, B**, Immunoblots for all 8 Cct subunits of the TRiC chaperonin complex (**A**) or various sHsps (**B**) for three independent samples of NSPCs and progeny cells (differentiated for 7 days (**A**) or 2, 4, and 7 days (**B**). Gapdh (**A**) and actin (**B**) serve as loading controls. **C**, Cct1 and Sox2 and Nestin (NSPC markers) in NSPCs. Scale bar: 10 µm. **D**, TRiC (***i***; Cct1 and Cct2 subunits) and HspB5 (***ii***), and Hoechst in NSPCs and differentiated cells. Traces: cell outlines. **E**, HspB5 with Dcx (neuronal marker) in *in vitro* differentiated cells (single channels in Figure S4G). **F**, Mouse brain sections of the SGZ (***i***) and SVZ (***ii***) NSPC niches, immunostained for Sox2 (NSPC marker) and TRiC (Cct1). Scale bars: 50 µm. Insets: Magnification of Sox2^+^/Cct1^+^ cells at indicated areas. **G**, Mouse SGZ brain sections immunostained for Sox2 (NSPC marker) and HspB5. Scale bar: 50 µm. Insets: Magnification of Sox2^+^/HspB5^−^ cells at indicated area.

Next, we compared the cellular distribution of TRiC and HspB5 in NSPCs and their progeny using immunofluorescence. We observed that the TRiC subunits Cct1 and Cct2 were highly abundant and diffusely localized in Sox2^+^/Nestin^+^ NSPCs (Figures 4C, D). Under these imaging conditions, TRiC was nearly undetectable in differentiated progeny. Of note, TRiC is essential and still present in differentiated cells, albeit at much reduced levels (e.g. Figure S4F for higher sensitivity imaging). In contrast to TRiC, HspB5 was undetectable in NSPCs but highly expressed in differentiated cells, including neurons, where it localized to a range of subcellular structures (from diffuse to punctate) (Figures 4D-E, S4G; as also previously reported (Golenhofen and Bartelt-Kirbach, 2015)).

The differential expression of these chaperone systems was next assessed in intact neural stem cell niches using immunohistochemistry. TRiC was strongly expressed in Sox2^+^, Pax6^+^, and Tbr2^+^ NSPCs in the SGZ and SVZ of the young adult mouse brain (Figures 4F; S4H, I), but expressed at much lower levels in surrounding cells (Figures 4F; S4H, Hi). Conversely, HspB5 was absent from Sox2^+^ SGZ NSPCs but distributed throughout the surrounding cells (Figure 4G). Of note, the constitutive Hsc70 isoform was uniformly distributed throughout the young adult SGZ niche, consistent with our data that its levels are not affected by differentiation (Figures 3, S4J). These findings demonstrate a switch in cellular chaperone networks during NSPC differentiation, from high TRiC and low sHSP expression in NSPCs to low TRiC and high sHSP levels in the differentiated cells. We next examined if these changes underlie their differential cellular proteostasis strategies to manage protein misfolding and toxicity.

### TRiC maintains NSPC proteome solubility

TRiC promotes *de novo* protein folding and is a potent suppressor of aggregation and toxicity of amyloidogenic proteins, including polyQ-expanded Htt and alpha-synuclein (Gestaut et al., 2019; Kitamura et al., 2006; Shahmoradian et al., 2013; Sontag et al., 2013; Tam et al., 2006). We considered whether the high TRiC levels contribute to suppressing aggregation in NSPCs. A time-course of TRiC expression during NSPC differentiation supported this idea. TRiC downregulation occurred early (within 24 h post-differentiation), together with NSPC-specific markers Sox2 and Nestin (Figure 5A*i*). TRiC was inversely correlated to the induction of the neuronal marker NeuN (Figure 5A*i*). Notably, we started to detect HttQ97 inclusions at 24 h post-differentiation, when Cct1 levels were reduced by over 50% (Figure 5A*ii*), suggesting TRiC may play a role maintaining solubility of misfolded proteins in NSPCs.

**Figure 5.**
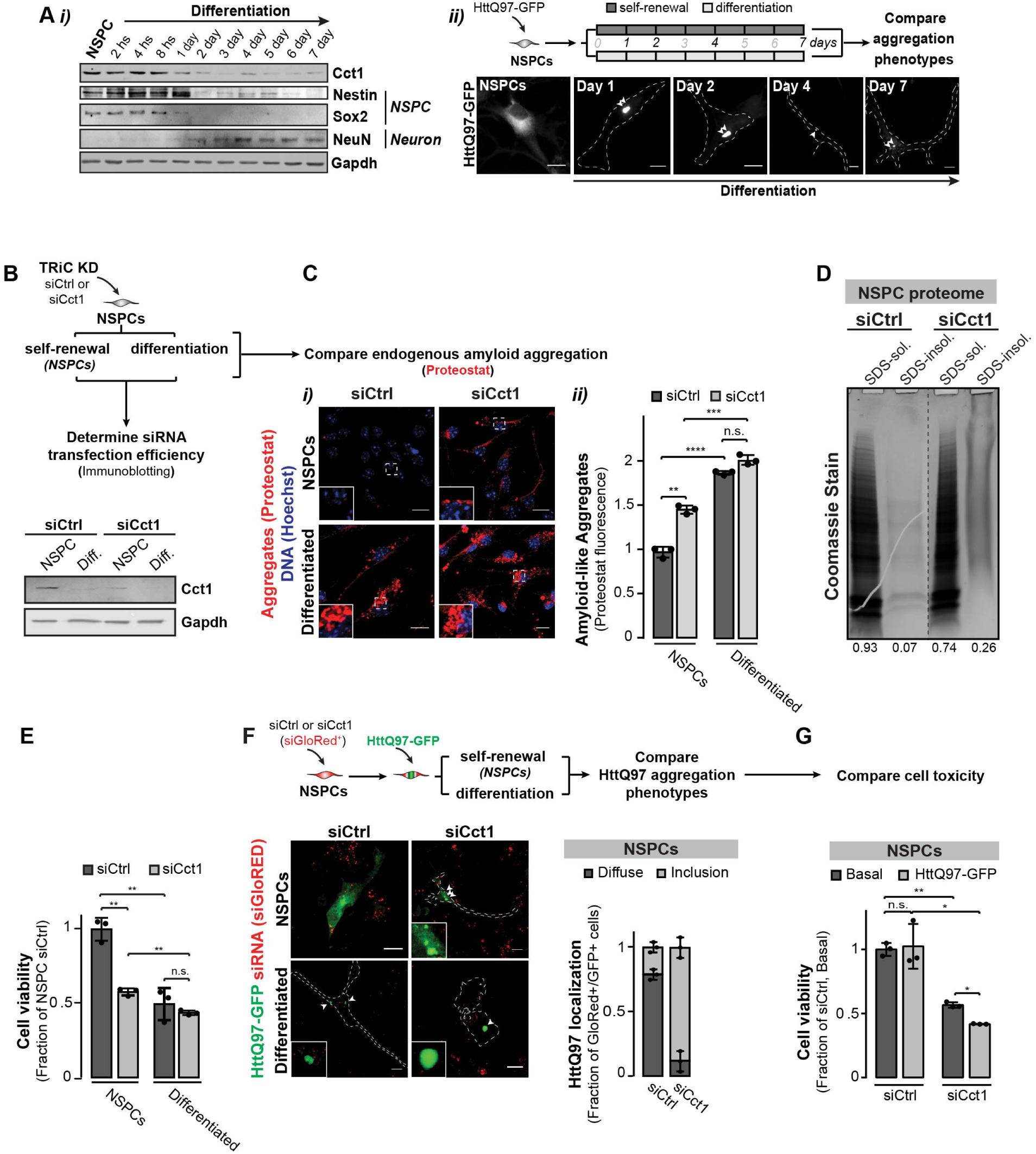
TRiC in NSPCs ensures proteome solubility and cellular fitness. **A**, (***i***) Immunoblot analyses of TRiC (Cct1), NSPC markers (Nestin, Sox2), and neuronal marker (NeuN). Gapdh serves as loading control. (***ii***) Epifluorescent images of HttQ97-GFP in NSPCs or during NSPC differentiation (at day 1, 2, 4, or 7). Traces: cell outlines; arrows: aggregates. **B**, Cct1 immunoblot of control (siCtrl)- and siCct1-targeted NSPCs as well as differentiated cells. Gapdh serves as loading control. Expression in siCtrl-treated NSPCs is set at 1. Degree of Cct1 knockdown is representative for all experiments presented. **C**, Aggregation of endogenous proteins in NSPC treated with control or a pool of siRNAs targeting *Cct1*; assessed by confocal imaging of Proteostat and DNA (Hoechst; (***i***). Insets: Magnification of Proteostat^+^ depositions at indicated areas); and quantified by fluorescence (mean ± SD; ***ii***). Proteostat fluorescence of siCtrl-targeted NSPCs is set at 1. ** p < 0.01; **** p < 0.001; n.s., non-significant. A similar increase in Proteostat^+^ aggregates was observed in NSPCs treated with single siRNAs targeting *Cct1* (Figure S5C) or a siCct2 SMARTPool (Figure S5F). **D**, siRNA-mediated Cct1 depletion from NSPCs impairs NSPC proteome solubility. Relative Coomassie signal, determined by densitometry, is displayed for each lane. **E**, MTT assay on control (siCtrl) and siCct1-treated NSPCs prior to or after differentiation (represented as mean ± SD). Metabolic activity of siCtrl-targeted NSPCs is set at 1. ** p < 0.01; n.s., non-significant. **F**, HttQ97-GFP and siGloRED (indicator of siRNA transfection) in control (siCtrl) and siCct1-treated NSPCs and differentiated cells. Arrows: aggregates. Insets: Magnification of HttQ97-GFP inclusions. Quantification (mean ± SD) of diffusely localized *vs*. aggregated HttQ97-GFP in double-transfected siCtrl/siGloRED^+^- and siCct1/siGloRED^+^-treated NSPCs (≥ 40 cells/condition/experiment). Similar results were obtained upon TRiC depletion using single siRNAs targeting *Cct1* (Figure S5D). **G**, Viability of control (siCtrl) and siCct1-treated NSPCs at basal conditions or upon exogenous expression of HttQ97-GFP (represented as mean ± SD). siCtrl-targeted NSPCs is set at 1. * p < 0.05; ** p < 0.01; n.s., non-significant. Similar results were obtained in NSPCs treated with a siCct2 SMARTPool (Figure S5G).

To directly test the role of TRiC in NSPC proteostasis, we employed a mixed siRNA pool (SMARTpool) that efficiently depletes cellular Cct1 and destabilizes the entire chaperonin complex (Freund et al., 2014) (Figures 5B, S5A). Co-transfection of the siRNAs with the fluorescent siGloRED siRNA transfection indicator confirmed we obtained comparable transfection efficiencies in NSPC and differentiated cell populations (Figure S5B). Immunoblot analysis of the transient siRNA-transfected population estimated an average TRiC depletion of ∼70 % over control (Figure 5B). TRiC depletion in NSPCs reduced the cellular proteome solubility, leading to significant increases in Proteostat^+^ aggregates of endogenous proteins, to a level approaching that of differentiated cells (Figures 5C, S5C). Direct biochemical analyses further indicated that Cct1 depletion from NSPCs reduced proteome solubility, leading to a marked increase in SDS-insoluble proteins (Figure 5D). Importantly, siRNA-mediated TRiC reduction in NSPCs significantly lowered the overall cellular fitness, to a level comparable to that of differentiated progeny (Figure 5E). Strikingly, TRiC depletion in NSPCs also reduced the diffuse HttQ97 localization and increased the fraction of cells with HttQ97 inclusions. TRiC depletion also sensitized NSPCs to HttQ97-GFP expression, suggesting this chaperonin prevents formation of toxic misfolded protein conformations (Figures 5F, G; S5D). Of note, similar results were obtained upon TRiC depletion with various single siRNAs against *Cct1* or multiple siRNAs targeting the *Cct2* subunit (Figures S5C-G). Together, these experiments support a central role for TRiC in the robust proteostasis capacity of NSPCs, and indicate that TRiC action precludes formation of toxic amyloids and HttQ97 aggregating species.

### HSPB5 promotes protein sequestration in differentiated progeny

We next examined the function of sHSPs induced in differentiated lineages. ATP-independent sHSPs are known to stabilize and protect misfolded proteins from irreversible aggregation by virtue of their alpha-crystallin domains (Haslbeck et al., 2019; Janowska et al., 2019). Paradoxically, increased sHSP levels during NSPC differentiation were accompanied by enhanced abundance of Proteostat-positive puncta and HttQ97 inclusions (Figures 1C; 2A, D). The sHSP from *S. cerevisiae* Hsp42 (herein yHsp42) also functions to promote active sequestration of misfolded proteins into spatially restricted protective inclusions termed Q-bodies, which enhance cellular fitness during stress (Escusa-Toret et al., 2013). Yeast cells lacking Hsp42 (*hsp42Δ*) fail to sequester misfolded proteins into inclusions, which reduces cellular fitness (Figure S6A, and (Escusa-Toret et al., 2013)). To test whether mammalian sHSPs may serve a similar spatial PQC function, we expressed human sHSPs in cells of the y*Hsp42* deletion strain and asked if they restore misfolded protein sequestration into inclusions. Significantly, several human sHSPs (i.e. HSPB3, HSPB5, HSPB6, and HSPB8) re-established Q-body formation in *hsp42Δ* cells (Figure S6A). These experiments indicate that some mammalian sHSPs induced during NSPC differentiation have the ability to promote spatial sequestration of misfolded proteins into PQC inclusions. Of note, HspB5, which promotes sequestration and is strongly induced upon NSPC differentiation, has been linked to HD pathology and associates with Alzheimer Disease’ fibers (Golenhofen and Bartelt-Kirbach, 2015; Oliveira et al., 2016).

To directly test the role of sHSPs in differentiated progeny, we focused on HspB5, which was depleted using pooled siRNAs (SMARTpool) as well as various single siRNAs (Figure S6B). Reducing HspB5 levels in differentiated cells led to a significant increase in their already high sensitivity to H_2_O_2_ treatment (Figure 6A). Strikingly, HspB5 depletion caused a dramatic decline in the fraction of differentiated cells with Proteostat^+^ puncta; suggesting it plays a role in inclusion formation (Figures 6B, S6C). We also examined how HspB5 affects HttQ97 management in differentiated cells. Consistent with a role promoting spatial sequestration, immunostaining indicated that HspB5 co-localizes with HttQ97-GFP inclusions (Figures 6C, S6D). HspB5 depletion reduced the fraction of cells with HttQ97-GFP inclusions compared to controls (siHspB5 *vs*. siCtrl; Figures 6D, S6E). The reduction in HttQ97 sequestered in inclusions in HspB5-depleted differentiated cells was accompanied by enhanced cell sensitivity to HttQ97 expression (Figure 6E). These results suggest that the formation of misfolded protein inclusions in differentiated cells is not a passive consequence of misfolded protein aggregation but rather a protective response mediated by sHSPs, which concentrate misfolded and disease-linked mutant proteins.

**Figure 6.**
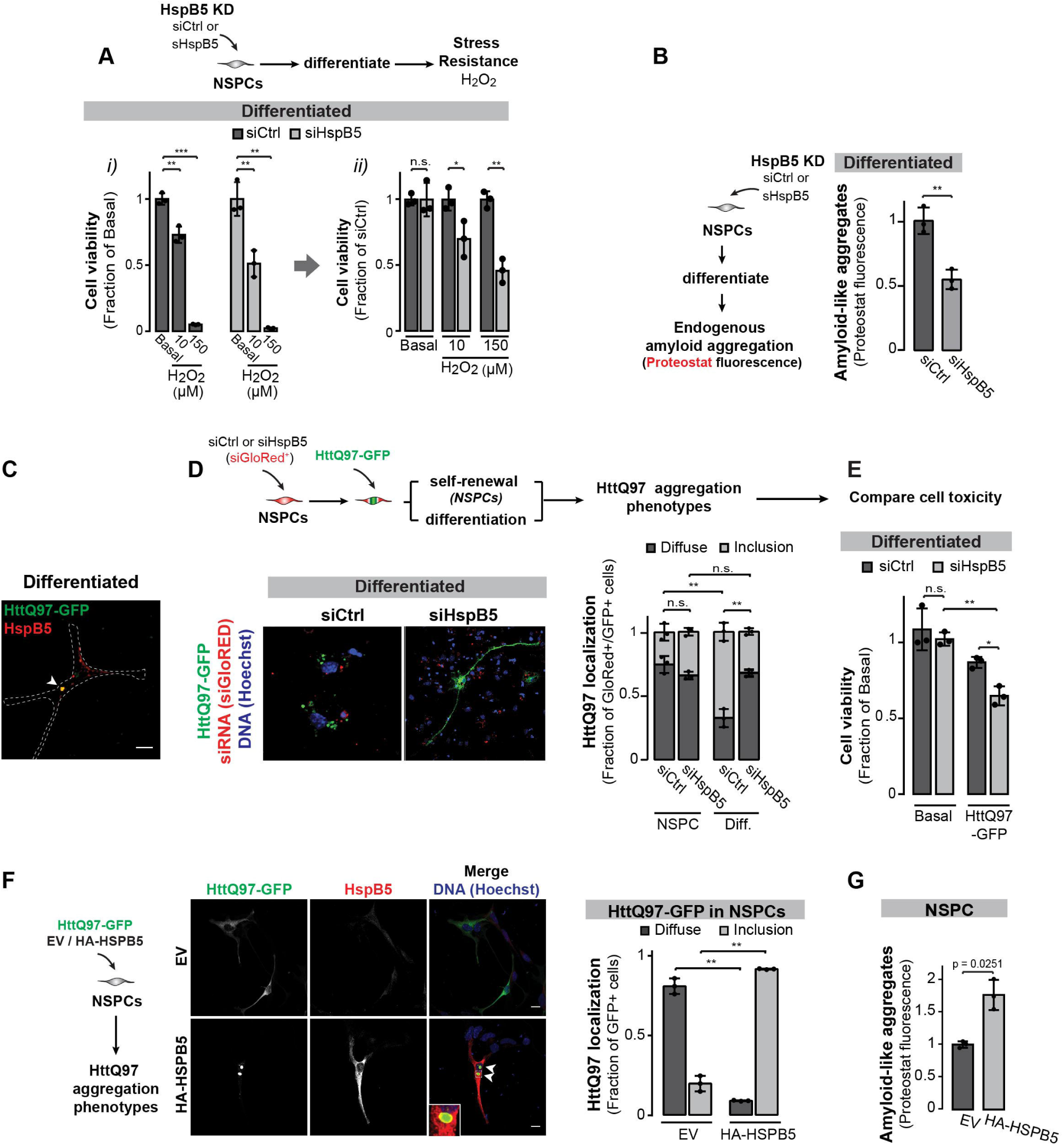
HSPB5 promotes sequestration of misfolded proteins in protective inclusions in differentiated cells. **A**, Relative viability (mean ± SD) of differentiated cells, treated with control siRNAs or a siHspB5 SMARTPool, at basal and oxidative stress (16 h, 10 or 150 μM H_2_O_2_) conditions. Viability of untreated cells for each condition is set at 1 (***i***). Viability of siCtrl- and siHspB5-treated cells at basal conditions is set at 1 (***ii***). * p < 0.05; ** p < 0.01; n.s. non-significant. **B**, Aggregation of endogenous proteins quantified by Proteostat fluorescence (mean ± SD) in differentiated progeny of control (siCtrl) and siHspB5-treated NSPCs. Fluorescence of siCtrl-targeted cells is set at 1. ** p < 0.01. Similar results were obtained upon HspB5 depletion using single siRNAs (Figure S6C). **C**, HspB5 and HttQ97-GFP inclusions in differentiated cells (single channels in Figure S6D). Trace: cell outline; arrow: HspB5 localization to HttQ97-GFP aggregate. **D**, HttQ97-GFP, siGloRED, and Hoechst in differentiated progeny of siCtrl- and siHspB5-treated NSPCs. Quantification (mean ± SD) shows diffusely localized *vs*. aggregated HttQ97-GFP (≥ 50 siRNA-targeted NSPCs and differentiated cells/experiment). Similar results were obtained in differentiated cells treated with various single siRNAs targeting HspB5 (Figure S6E). **E**, Relative viability (mean ± SD) of control (siCtrl) and siHspB5-treated differentiated cells under basal and proteotoxic stress (exogenous HttQ97-GFP) conditions. Viability of siCtrl-treated cells for each condition is set at 1. * p < 0.05; ** p < 0.01; n.s. non-significant. **F**, NSPCs exogenously expressing HttQ97-GFP and EV or HA-HSPB5. DNA stained with Hoechst. Arrows: HttQ97-GFP inclusions. Inset: Magnification of HA-HSPB5 co-localization at HttQ97-GFP inclusion at indicated area. Quantification (mean ± SD) shows diffuse *vs*. aggregated HttQ97-GFP (≥ 50 NSPCs/condition/experiment). **G**, Fluorescent quantification (mean ± SD) of amyloid-like aggregates in control (EV) and HA-HSPB5 expressing NSPCs. p = 0.0251.

As a test of this hypothesis, we ectopically expressed HA-tagged HSPB5 in NSPCs, which do not form inclusions and have high levels of TRiC. Remarkably, HSPB5 expression promoted sequestration of HttQ97 in inclusions that partially co-localized with HA-HSPB5 (Figure 6F). In addition, we also observed the emergence of endogenous Proteostat^+^ inclusions (HA-HSPB5 *vs*. EV transfected cells; Figure 6G). These data indicate that sHSPs promote spatial misfolded protein sequestration, even in the presence of a potent soluble chaperone machinery. This supports the notion that sHSPs induction in neural progeny cells serves to switch the cellular proteostasis strategy to spatial sequestration of misfolded proteins into inclusions to enhance cellular fitness.

### TRiC levels decline during aging

We next examined how aging affects TRiC and sHSPs levels in the brain. Proteostat staining of aged brain sections compared to young counterparts indicated that the increase in Proteostat^+^ inclusions was not restricted to aged NSPCs (Figures 1D-G, S1C), but was widespread in other regions; particularly the cortex and striatum (Figures 7A, B; S7A), areas typically affected in age-related proteopathies (Kalin et al., 2017; Rub et al., 2016). We observed a decline in TRiC protein levels in Sox2^+^ cells of the aged SGZ stem cell niche compared to those of young adult mice; both *in vivo* in mouse brain sections (Figures 7C, S7B) and *in vitro* in isolated NSPCs (Figures 7D). Of note, aging also reduces TRiC expression in other brain areas, including the striatum and cortex (Figures 7C*ii*, S7B). Analysis of *Cct1* mRNA expression in aged NSPCs was significantly reduced compared to young NSPCs (Figures 7E, S7C); but for other subunits the age-dependent reductions were more variable (Figure S7C). Thus, the age-linked decline in TRiC expression may not only be driven by transcriptional changes, although lower levels of one CCT subunit will reduce assembly of the entire TRiC complex (Freund et al., 2014). In contrast to TRiC, HspB5 levels exhibited no significant aging-linked alteration in mouse brains; though minor induction of a few other sHsps was detected (Figures S7D-F). Thus, these findings are indicative of an aging-related decline in NSPC proteostasis, particularly TRiC, contributing to the onset and progression of neurodegenerative proteopathies.

**Figure 7.**
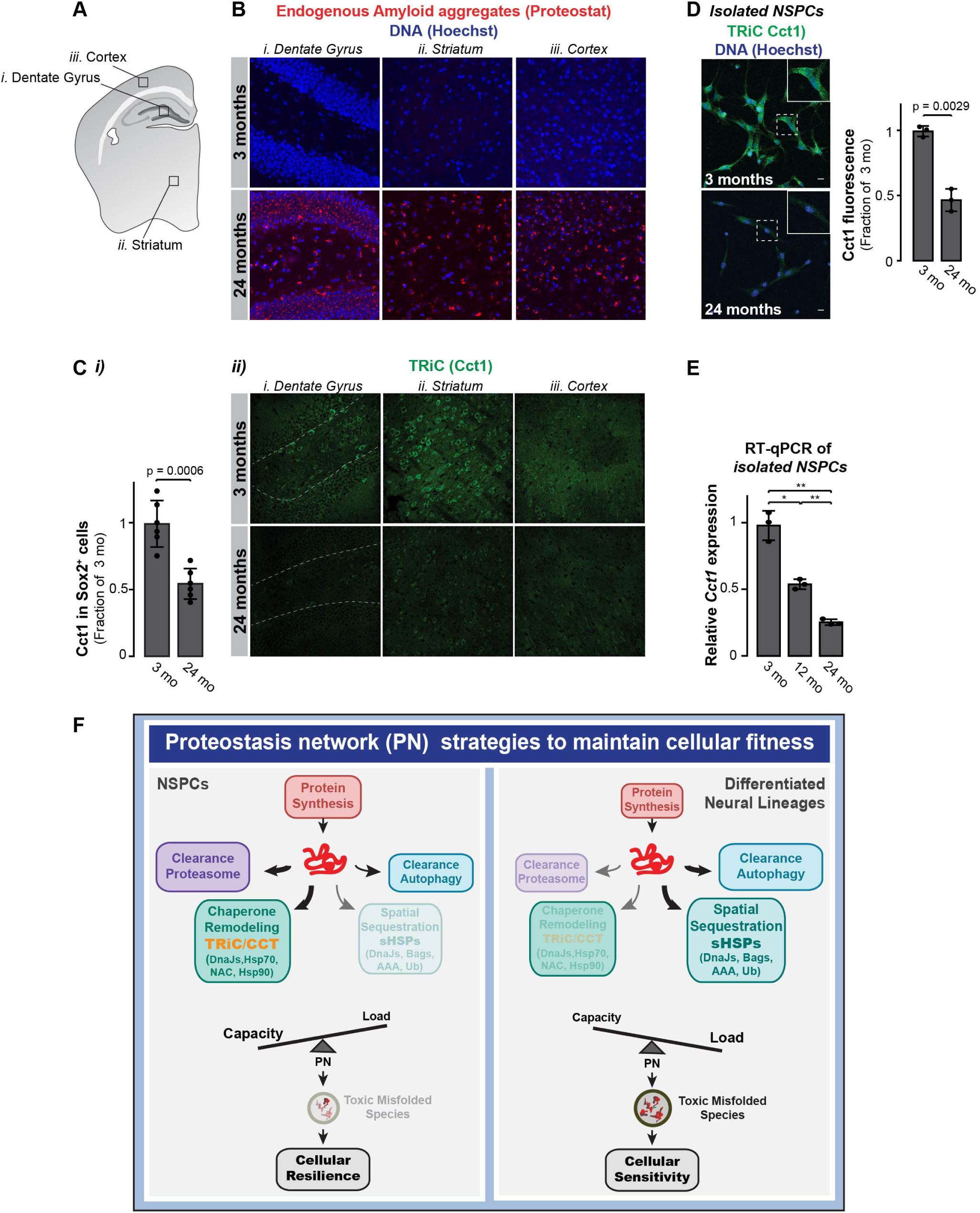
Aging attenuates TRiC expression in NSPCs as well as the overall brain. **A**, Schematic overview of coronal brain section. Areas of images shown in **B** and **C** are indicated. **B**, Proteostat^+^ amyloid deposits in brain sections from young and old mice in Dentate Gyrus, Striatum, and Cortex. See Figure S7A for stitched images of full brain sections. **C**, (***i***) Quantification (mean ± SD) of Cct1 fluorescence in Sox2^+^ cells (whole frame; 6 mice/age group). Cct1 fluorescence in young mouse brain is set at 1. p = 0.0006. See Figure 7B for stitched images of Cct1/Sox2/Neun stained brain sections; (***ii***) Cct1 in brain sections from young and old mice in Dentate Gyrus, Striatum, and Cortex. **D**, TRiC (Cct1) and Hoechst in young and old isolated NSPCs. Insets: Magnification of cellular Cct1. Quantification (mean ± SD) shows Cct1 fluorescence in young and old NSPCs. Fluorescence of young NSPCs is set at 1. p = 0.0029. **E**, *Cct1* expression (mean ± SD) in young, middle-aged, and old NSPCs. mRNA abundance in young NSPCs is set at 1. * p < 0.05; ** p < 0.01. **F**, Differentiation drastically remodels the robust chaperone network and proteostasis logic of NSPCs to enhance cellular fitness. NSPCs express high levels of TRiC, which is central to maintain a soluble proteome and contributes to the NSPC ability to effective manage non-native proteins and withstand proteotoxic stress. Differentiation drastically remodels the proteostasis logic. In contrast to NSPCs, the load of misfolded proteins is markedly increased in differentiated cells; most likely due to a marked attenuation of TRiC expression, protein synthesis rates, and UPS clearance activities. sHSPs are induced during NSPC differentiation. This promotes spatial sequestration of misfolded proteins into defined protective inclusions to clear the cellular milieu of potentially harmful, misfolded proteins.

## Discussion

The ability of adult NSPCs to self-renew and differentiate into multiple neural lineages makes them indispensable for cognitive plasticity and cellular replacement, e.g. during neurodegeneration. We find NSPCs differentiation involves a shift in the logic of proteome maintenance through differential expression of core chaperone machineries with distinct modes of action, which guide non-native proteins to distinct cellular fates (Figure 3F).

Eukaryotic cells contain distinct chaperone classes guiding non-native polypeptides to their folded state or promoting PQC through refolding, degradation, and sequestration (Figure 3F). While all chaperones share the ability to recognize non-native polypeptides, they differ in their structural and mechanistic characteristics as well as in their molecular interactions within cells, including those with other proteostasis factors. NSPCs express high levels of ATP-dependent chaperonin TRiC, central to maintaining a soluble proteome and to the cellular ability to manage mutant proteins and withstand proteotoxic stress (Figure 7F). TRiC binds nascent and non-native polypeptides in its central chamber, preventing aggregation and promoting either protein folding or clearance. By contrast, differentiated neural cells strongly downregulate TRiC while inducing sHSPs; ATP-independent chaperones that stabilize non-native proteins and promote their spatial sequestration into protective inclusions (Figure 7F). Our finding that human sHSPs can, by themselves, promote misfolded protein sequestration into inclusions in yeast indicates this is an autonomous activity of this class of chaperones. Furthermore, ectopic HspB5expression in NSPCs promotes formation of HttQ97 and endogenous amyloid inclusions. We speculate that the intrinsically disordered domains in sHSPs allow them to form phase-separated puncta that clear the cellular milieu of potentially harmful, misfolded proteins. Indeed, downregulation of HspB5 in differentiated cells increases their vulnerability to proteotoxic insults.

Our experiments revealed a central role played by changes in TRiC and sHSPs levels in a switch in proteostasis logic upon NSPCs differentiation. However, it is important to note our analysis also identifies rewiring of other chaperone co-factors and PQC factors (Figure 3). For instance, differentiation downregulates subunits of the co-translationally acting chaperone NAC (i.e. NACA and BTF3), which, similar to TRiC, also plays a role suppressing aggregation (Koplin et al., 2010; Rospert et al., 2002; Shen et al., 2019). Further, we observed differentiation-related changes in Hsp70/Hsp90 co-factors, including DnaJ and Bag proteins, which act to recruit these Hsp70/Hsp90 chaperones to various cellular structures and factors (Kampinga and Craig, 2010; Taguwa et al., 2015). Future studies should explore how these changed chaperone networks interface with either TRiC or sHSPs to promote cell-specific proteostasis. Overall, these changes illustrate how chaperone networks can be developmentally reshuffled to provide a proteostasis strategy that suits the biological needs of different cell types.

It is intriguing to consider the functional or metabolic constraints driving the changed proteostasis strategy. In principle, the high translation and cell division rates in NSPCs may require high TRiC levels (Gestaut et al., 2019), while the extensive cytoskeletal network of neurons may require stabilization by sHSPs (Singh et al., 2007). A potential rationale for the shift in proteostasis strategy may be the balance between metabolism and the demands on the energetic supply of a cell. TRiC is an energy-intensive chaperone, imposing an energetic cost to the high protein solubility and high stress resilience of NSPCs. The high levels of translation, DNA replication, and proteasome activity in NSPCs also contribute to an energetically demanding physiology. The need to maintain the NSPC regenerative potential throughout life may justify a more stringent PQC system, while differentiated cells may adequately manage their proteome with a less ATP-intensive system. The different logic and composition of the cellular chaperone networks may thus reflect a tradeoff between the physiological function and energetic demands of a given cell.

Understanding chaperone networks in distinct neural lineages may illuminate their vulnerability to amyloid diseases, particularly those linked to late-onset neurodegeneration. Based on our findings (Figures 1D-H, 7A-E), we speculate that aging generally reduces the levels of TRiC and likely other chaperone systems. Indeed, TRiC subunit downregulation was recently reported in striatal medium spiny neurons generated from HD patient fibroblasts at pre-symptomatic and symptomatic disease stages and found to be correlated with cellular age (Victor et al., 2018). In further agreement with our findings, longitudinal microarray analyses of human brain postmortem samples also find TRiC levels decline in aged human brains. Given TRiC’s anti-aggregation functions, such a decline may contribute to late-onset aggregation-diseases (Brehme et al., 2014). Thus, the mechanisms of the aging-related decline in NSPC proteostasis may mirror those observed in other tissues. Alternatively, they may reflect age-related changes in the NSPC pool, such as increased quiescence due to neuro-inflammatory signals (Doetsch et al., 1999; Morshead et al., 1994; Seri et al., 2001). Interestingly, quiescent NSCs have lower TRiC transcript levels and accumulate large SDS-insoluble protein inclusions (Leeman et al., 2018).

The observation that NSPCs and differentiated progeny employ different proteostasis strategies to manage misfolded proteins helps rationalize disparate sets of previous observations regarding the link between Htt inclusion formation and toxicity. The high levels of TRiC in NSPCs explain studies in an HD mouse model showing that NPCs in the SVZ were almost completely devoid of Htt inclusions; which only appeared as upon differentiation into neurons (Benraiss et al., 2013). Our finding that high sHSP levels mediate protein sequestration in differentiated neural lineages may explain why experiments in post-mitotic neurons found sequestration of HttQ97 into inclusions to be protective and promoting survival (Arrasate and Finkbeiner, 2012; Arrasate et al., 2004). Our findings provide a conceptual framework to interpret these observations. High TRiC levels in NSPCs maintain aggregation-prone HttQ97 in a soluble but non-toxic state. In these cells, the visible inclusions following decreased proteostasis capacity accompany proteotoxicity. By contrast, differentiated neural progeny detoxify misfolded proteins through sHSP-mediated sequestration into protective inclusions. We hypothesize that toxic and protective inclusions will differ fundamentally in their genesis, organization, and composition, and their characterization will require imaging by super-resolution microscopy (Sahl et al., 2016; Sahl and Vonk, 2019) or cryo-electron tomography (Bauerlein et al., 2017). A better understanding of the logic of misfolded protein management in affected cells is instrumental to inform strategies to prevent or delay proteotoxic diseases.

## Supporting information

Table S2. Differential gene expression between NSPCs and differentiated neural progeny cells

Table S3. REACTOME enriched terms in NSPCs vs. differentiated neural progeny cells (

## Acknowledgments

We gratefully acknowledge the National Institute for Aging (NIA), Dr. J. Middeldorp (University Medical Center Utrecht, Utrecht, the Netherlands), and Prof. T. Wyss-Coray (Stanford University, CA, USA) for kindly providing us mice from their aging colony and paraformaldehyde-perfused brain sections of young and aged mice. We thank J. Rothbard (Stanford University, Stanford, CA, USA) and Prof. H. H. Kampinga (University Medical Center Groningen, Groningen, the Netherlands) for providing the sHSPB1-B9:pET21 and AR-Q113-GFP constructs, respectively. We thank members of the Frydman and Brunet labs and Martijn Gloerich for useful comments and discussions.

This work was supported by NIA grant no. AG056290 (JF/AB), the Marie Curie International Outgoing Fellowship Program (to WIMV; 303096 FOLDEG), NIH/NIGMS National Research Service Award (to TKR, F32GM120947), the Glenn Laboratories Seed Grant Program, and the CHDI Foundation (JF).

## Author contributions

Conceptualization, WIMV and JF; Methodology, WIMV, AEW, and AB; Software, PTD; Validation, WIMV and TKR; Formal Analysis, WIMV, TKR, and PTD; Investigation, WIMV, TKR, PTD, and AEW; Resources, WIMV, AB, and JF; Data curation, PTD; Writing – Original draft, WIMV and JF; Writing – Review & Editing, WIMV, TKR, PTD, AEW, AB and JF; Visualization, WIMV and JF; Supervision, JF; Project Administration, WIMV and JF; Funding Acquisition, WIMV, TKR, AB and JF.

## STAR Methods

## LEAD CONTACT AND MATERIALS AVAILABILITY

Further information and requests for resources and reagents should be directed to and will be fulfilled by the Lead Contact, Judith Frydman.

## EXPERIMENTAL MODEL AND SUBJECT DETAILS

### Animals

Wild-type FVB/N female mice were purchased from Jackson Laboratories, and C57/Bl6 female mice for the aging experiments were kindly provided by the NIA aging colony. All care and procedures were in accordance with the Guide for the Care and Use of Laboratory Animals. All animal experiments were approved by Stanford’s Administrative Panel on Laboratory Care (APLAC) and in accordance with institutional and national guidelines. The animals were housed under standard laboratory conditions on a 12 h light/dark cycle with constant access to food and water.

### Mouse NSPC Isolation

Isolation of adult mouse NSPCs from brain tissue was performed as described previously (Palmer et al., 1997). In brief, the brains were dissected from five-seven 3 month-old mice and the olfactory bulbs, cerebellum, and brainstem were removed. The forebrains were finely minced, digested for 30 min at 37 °C in HBSS media containing 2.5 U/ml papain (Worthington), 1 U/ml Dispase II (Roche), and 250 U/l DNase I (Sigma), and mechanically dissociated. Neural stem cells and progenitor cells were purified with two Percoll gradients (25 % followed by 65 %; Amersham), and plated at a density of 10^5^ cells/cm^2^ in self-renewal media (i.e. NeuroBasal-A medium (Invitrogen) supplemented with antibiotics (penicillin-streptomycin-glutamine; Invitrogen), 2 % B27 supplement without vitamin A (Invitrogen), and growth factors bFGF and EGF (both 20 ng/ml; Peprotech). For aging experiments, NSPCs were isolated from brains dissected from 3 (young adult), 12 (middle-aged), or 24 (aged) month old mice.

### NSPC culture and differentiation

Cells were incubated at 37 °C in 5 % CO_2_ and 20 % oxygen at 95 % humidity. Media was replaced every other day. For *in vitro* experiments, isolated NSPCs were expanded as neurospheres *in vitro* in self-renewal media as described in Renault *et al*. (Renault et al., 2009). Neurospheres were dissociated with Accutase (Chemicon), plated as single cells on poly-D-lysine (Sigma) coated plates or glass coverslips, and grown as proliferating cultures in self-renewal media. NSPC differentiation was induced (in general, one day after plating) by replacing the self-renewal media with differentiation media (NeuroBasalA medium supplemented with penicillin-streptomycin-glutamine, 2 % B27 supplement without vitamin A, and 0.5 % fetal bovine serum (FBS; Gibco)) following a gentle one-time wash with differentiation media to remove growth factor residues. Cells were incubated at 37 °C in 5 % CO_2_ and 20 % oxygen at 95 % humidity. Media was replaced every other day. Unless stated otherwise, NSPCs and differentiated cells were fixed/harvested and processed in parallel. The majority of experiments presented represent NSPCs at passage number 3-6 (NSPC cultures were discarded at passage 8). Notably, no significant variations between NSPC passages were observed (see Figure S4B). Unless stated otherwise, cells were differentiated for seven days prior to analyses.

### Free-floating brain sections

Free-floating brain sections (30 µm) of young adult (3 month old), middle-aged (12 month old), and aged (24 month old) mice were either kindly provided by Dr. J. Middeldorp (University Medical Center Utrecht, Utrecht, the Netherlands), and Prof. T. Wyss-Coray (Stanford University, CA, USA).

## METHOD DETAILS

### Plasmids and strains

For mammalian expression, HttQ97-GFP:pcDNA3,1 and VHL^L158P^-GFP:pcDNA3,1 were described previously (Kaganovich et al., 2008). The N-end rule and ubiquitin fusion degradation (UFD) reporter plasmids used to express short-lived green fluorescent proteins for quantification of ubiquitin-proteasome-dependent proteolysis (i.e. Ub-M-eGFP (stable), Ub-R-eGFP (N-end rule degradation signal), and Ub-G76V-eGFP (UFD signal); all three in eGFP-N1 plasmid backbone) were previously described by Dantuma *et al*. (Dantuma et al., 2000) and commercially obtained from Addgene plasmid repository. SOD1^G37R^-GFP was generated by site-directed mutagenesis on the SOD1^WT^-GFP:pcDNA3.1 plasmid (Vonk et al., 2014). ARQ113-GFP was kindly provided by H.H. Kampinga (UMCG, Groningen, the Netherlands). The *Tcp1*-Gaussia Luciferase (*Tcp1*-GLuc) GLuc-ON promotor clone containing a ∼ 1 kb fragment upstream of the transcription start site and a ∼ 0.3 kb fragment of the exon 1 of the murine *Tcp1* in pEZX-PG04 vector. This bi-cistronic vector also constitutively expresses a secondary reporter, secreted alkaline phosphatase (SEAP), which is used as internal transfection control (GeneCopoeia).

For yeast expression, the BY4741 wild-type strain and the *hsp42*Δ strain, and Ubc9-2:pAG416Gal1-GFP and yHsp42:pRS313GPD constructs were described previously (Escusa-Toret et al., 2013). Untagged human sHSPB1-B9:pET21 constructs were kindly provided by J. Rothbard (Stanford University, Stanford, CA, USA), and subcloned into the pAG426GPD-eGFP-ccdB Gateway plasmid using EcoRI and HindIII restriction sites.

### Reagents

The primary and secondary antibodies for SDS-PAGE Western blot analysis (WB), immunohistochemistry (IHC), and immunocytochemistry (ICC) are listed in STAR Methods Key Resources Table. DNA was stained with Hoechst 33342 (Life Technologies).

Indicator of siRNA-transfection (siGloRED) as well as non-targeting pools of siRNAs (siCtrl) and single and pools of multiple siRNA (SMARTpool) targeting mCct1, mCct2, and mHspB5 were purchased as ON-TARGETplus siRNAs from Dharmacon.

### Cell treatments and transfection

To impair protein clearance by the proteasome, 3-15 µM MG132 (Z-Leu-Leu-Leu-al, Sigma) was added to the media for the indicated hours prior to cell lysis/fixation. Vehicle control (DMSO) treated samples were taken along in every experiment. To impair the proliferation rate of NSPCs, cells were treated with mitomycin C (MMC; Sigma) at indicated concentrations for indicated hours prior to fixation.

Cells were transfected using Lipofectamine2000 (Life Technologies), according to the manufacturer’s protocol, in a 1:3 DNA:Lipofectamine2000 ratio. Unless stated otherwise, per experiments, NSPCs were transfected using a transfection master mix. Four hours post-transfection, media was replaced and samples were split and cultured in either self-renewal or differentiation media to generate a NSPC and differentiated progeny pool.

siRNAs were transfected according to manufacturer’s protocol, using DharmaFECT as transfection agent. Media was replaced 18 h post-transfection. siRNA-treated cells were analyzed four days post-transfection. For some experiment, siRNA-transfection was followed by another round of transfection using Lipofectamine to introduce HttQ97-GFP into the cells (in general, one day post-siRNA transfection). In these experiment, only double-transfected cells (siGloRED^+^/HttQ97-GFP^+^) were included in the analyses. Unless stated otherwise, NSPCs were treated with siRNAs and maintained under self-renewal or differentiation conditions prior to analysis.

### RNA isolation and quantitative reverse transcription PCR (RT-qPCR)

Total RNA was extracted using TRIZOL^®^/Chlorophorm/Phenol method. RNA was converted to cDNA using the iScript reverse transcriptase kit (BioRad) following manufacturer’s instructions. Gene expression was detected using SybrGreen RT-qPCR expression assays in a 96-wells plate format as described by the manufacturer using MyIQ real-time PCR cycler (BioRad); cDNA templates were used at a 1:40 dilution. RT-qPCR primer sequences for all genes analyzed in this study are listed in Table S4.

### RNA-Sequencing

Cultures of NSPCs and differentiated progeny (three independent isolates, each at passage 3) were washed twice in cold PBS, and total RNA was purified using the QuickRNA Miniprep kit (Zymo Research). Samples were submitted to the Stanford Protein and Nucleic Acid Facility for quality assessment by Agilent Bioanalyzer to determine the RNA integrity (RIN) for each sample, which was > 9. Libraries were prepared using the TruSeq Stranded Total RNA Library Prep Gold Kit (Illumina) starting with ≥ 100 ng of total RNA from each sample. Libraries were quantified by KAPA Library Quantification Kit (Roche), normalized, and pooled for sequencing. All libraries were sequenced for High Output on an Illumina NextSeq 550. Sequencing was performed at the CZ Biohub facility at Stanford University. To characterize differential expression of pathways involved in differentiation and the change in proteostasis, we used a resampling method to estimate the significance of differential expression in REACTOME pathways (Croft et al., 2014).

### Proteasome fluorogenic chymotrypsin-like activity assay

The *in vitro* assay of biochemical measurement of chymotrypsin-like proteasomal activity was performed as described previously (Vilchez et al., 2012). In brief, cells were collected in proteasome activity assay buffer (50 mM Tris-HCl, pH 7.5, 250 mM sucrose, 5 mM MgCl_2_, 0.5 mM EDTA, 2 mM ATP, and 1 mM dithiothreitol), and lysed by passing ten times through an insulin syringe prior to centrifugation at 10,000*g* for 10 min at 4 °C. Protein concentrations were determined by BCA protein assay (Pierce). 15 µg of total protein (cell lysates) were transferred to a black 96-well microtiter plate (Corning) following addition of the proteasome substrate (Suc-Leu-Leu-Val-Tyr-AMC), which becomes highly fluorescent upon proteolytic cleavage. Incubation with MG132 was taken along to inactivate proteasome activity (negative control), whereas other cells were treated with DMSO as vehicle control. Fluorescence (Ex: 360 nm, Em: 480 nm) was monitored on a microplate fluorimeter (Infinite M1000, Tecan) every 5 min for 60 min at 37 °C. For each experiment, AMC fluorescent signal of each sample was normalized to the median AMC fluorescence intensity of all samples in the experiment.

### Western blot analysis

For protein analysis, cells were lysed in protein lysis buffer (10 mM Tris-HCl, pH 7.4, 10 mM EDTA, 50 mM NaCl, 1% Triton X-100) supplemented with 2 mM sodium orthovanadate, 1 mM phenylmethyl-sulphonyl fluoride, and Complete mini protease inhibitor cocktail mix (Roche) for 10 min at 4 °C, and cleared by centrifugation at 16,000*g* for 10 min at 4 °C. Protein concentrations were determined by BCA protein assay (Pierce), and 30 µg total protein lysate was used for SDS-PAGE Western blot analyses. Samples were denatured at 95 °C for 5 min. Proteins were transferred to 0.45 μm Immobilon-FL PVDF membranes (Millipore), and membranes were blocked in either 5 % BSA or Odyssey Blocking buffer (LI-COR Biosciences) followed by primary and secondary antibody incubations (IRDye 800CW or 680RD secondary (LI-COR Biosciences) antibodies were used). Uncropped images of all immunoblots presented are provided in the Supplemental Data File.

### Viability assays

Unless stated otherwise, cell viability was measured by MTT assay. NSPCs were seeded in a 96-wells plate at 10,000 and 20,000 cells/well in self-renewal media (four replicates per sample condition), and media was replaced the next day or following transfection with self-renewal (wells with 10,000 cells) or differentiation (wells with 20,000 cells) media. In some experiments, cells were exposed to oxidative (various concentrations H_2_O_2_ for 16 h at 37 °C) or heat (6 h at 42 °C) stress prior to media removal, warm PBS wash, and 5 min incubation (at 37 °C) of Hoechst dye (Life Technologies; 1:15,000 dilution) in warm PBS. Following warm PBS wash, the total number of cells/well were calculated by Hoechst fluorescence staining of nuclei using a fluorimeter (Tecan, 37 °C; Ex: 350 nm, Em: 461 nm). Next, PBS was replaced by NeuroBasalA-medium supplemented with MTT (0.4 mg/ml 3-(4,5-Dimethylthiazol-2-yl)-2,5-diphenyltetrazolium bromide, 98%; Sigma), and cells were incubated for 30-60 min in a 37 °C incubator. NeuroBasalA-medium (no MTT addition) – incubated cells were taken along as negative control for background subtraction. Cells were washed with PBS and lysed with isopropanol/0.04 M HCl. The quantity of formazan produced (directly proportional to the number of metabolically active, viable cells) was measured in a spectrophotometer (Em: 530 nm, Ex: 595 nm).

For quantification of cell death by fluorescence-activated cell sorting (FACS) analysis using the fluorescent, cell-impermeant dye 7-amino-actinomycin D (7AAD, BioLegend), cells grown in 6-wells plates were collected upon trypsinization, counted, and cell pellets were resuspended in 0.5 ml of cell staining buffer (BioLegend) supplemented with 5 µl of 7AAD per million cells. Samples (10,000 events per sample) were analyzed after 5-10 min incubation in the dark on ice on a FACSCalibur flow cytometer (BD Biosciences) in the FL3 channel.

### Immunocytochemistry

NSPCs were grown as adherent cells on coverslips pre-coated with poly-D-lysine. Prior to fixation with 4% paraformaldehyde for 15 min, cells were washed once with ice-cold PBS. Subsequently, cells were quenched with 10 mM NH_4_Cl for 5 min at room temperature following permeabilization (three times wash with 0.2% Triton X-100 in PBS) and blocking using blocking buffer (5% normal donkey serum (NDS) in PBS, 0.45 µm filtered) for 1 h at room temperature in the dark. Immunolabeling was performed in blocking buffer with the indicated primary antibodies for 2 h at room temperature in the dark. After three times washing in PBS/0.1% Triton X-100, secondary labelling was performed with secondary antibodies (diluted 1:500 in blocking buffer) for 1 h at room temperature in the dark. Slides were washed five times with PBS preceding mounting of coverslips in Immumount (ThermoScientific). Hoechst staining was performed in the one before final wash step.

### Immunohistochemistry

Free-floating brain sections were washed three times in TBS in a 12-well transwell plate for 5 min on a shaking platform prior to antigen retrieval (30 min incubation with preheated sodium citrate, pH 6.0 / 0.05% Triton X-100 at 80 °C). Sections in solution were cooled down for 20 min and washed three times in TBS following blocking in blocking buffer (TBS supplemented with 1% NDS, 1% BSA, 0.3% Triton X-100, and 0.01% sodium azide) at room temperature for 2 h on a shaking platform in a 48-well plate format. Primary antibodies were diluted in blocking buffer and labelling was done overnight (with agitation) in blocking buffer in a 96-wells plate format. Following three 5 min washes with TBS, sections were incubated for 2 h at room temperature with secondary antibodies (at a dilution of 1:400 in blocking buffer) in the dark on a shaking platform. Sections were rinsed three times for 5 min with TBS, and mounted using Vectashield with DAPI stain. For Proteostat staining of brain sections, sections were rinsed with PBS and incubated with Proteostat dye diluted in PBS according to manufacturer’s protocol. Sections were washed with PBS and destained with 1% acetic acid in PBS for 20 min on a shaking platform at room temperature. Sections were rinsed three times in PBS and either mounted in Immumount, or, when additional immunostainings were included, incubated for 2 h with blocking buffer prior to immunostaining (performed as described above).

### Quantitative amyloid-aggregation assay

NSPCs were seeded as adherent cells in a poly-D-lysine coated 96-wells plate at 10,000 and 20,000 cells/well in self-renewal media (four replicates per sample condition) and media was replaced the next day or following transfection with self-renewal (wells with 10,000 cells) or differentiation (wells with 20,000 cells) media. Prior to fixation with 4% paraformaldehyde for 15 min, cells were washed once with ice-cold PBS. Proteostat and Hoechst stainings were done according to manual instructions (Enzo Life Sciences), and fluorescence was measured in a Tecan fluorimeter (Proteostat: Ex: 550 nm, Em: 600 nm; Hoechst: Ex: 350 nm, Em: 461 nm).

### Fractionation of SDS-soluble and -insoluble fractions

Determination of NSPC proteome solubility was done according to the fractionation protocol reported by Juenemann *et al*. (Juenemann et al., 2015). Briefly, prior to collection and centrifugation of cells at 2,000 rpm for 5 min (4 °C), NSPCs, seeded as adherent cells in a poly-D-lysine coated 6-well plate at 250,000 cells/well in self-renewal media and as indicated treated with siRNAs for 48-72h, were washed once in cold PBS. Cell pellets were lysed in TEX buffer (70 mM Tris-HCl, pH 6.8, 1.5 % SDS, 20% glycerol). Upon brief vortexing, DNA was sheared by passing the cell lysate six times through a 21 Gauge needle. Next, proteins were reduced by addition of 1 M DTT (freshly made), and samples were boiled for 10 min at 99 °C in a thermoblock while shaking at 1,000 rpm prior to centrifugation for at least 60 min at 14,000 rpm (RT). The supernatant, representing the SDS-soluble fraction, was carefully collected. The pellet fraction containing the SDS-insoluble protein complexes were completely solubilized in 100 % formic acid and incubation at 37 °C for 40 min while shaking in a thermoblock at 1,000 rpm. Please note that the formic acid-treatment of lysates solubilizes aggregation-prone mHTT for SDS-PAGE analysis (Hazeki et al., 2000). The formic acid was evaporated using a Speedvac. The remaining pellet was dissolved in 50 μl TEX buffer supplemented with a bromophenol blue. Prior to loading on a 4-20% mini-PROTEAN TGX precast protein gel (BioRad), samples were boiled for 10 min at 99 °C in a thermoblock while shaking at 1,000 rpm. The SDS-soluble and SDS-insoluble fractions were loaded in a 1:5 ratio. For visualization of NSPC proteome by Coomassie staining, gels were fixed for 4 h in 50 % Methanol/10 % gliacal acetic acid (fixation solution was changed once an hour). Subsequently, the gels were stained for 20 min in 0.1 % Coomassie R250 / 50 % Methanol / 10 % glacial acetic acid followed by destaining in 40 % Methanol. Gels were stored in 5% glacial acetic acid prior to imaging.

### Luciferase-based promoter-activity assay

NSPCs were plated onto 96-well plates (10,000 and 20,000 cells/well, respectively) one day prior to transfection with the Tcp1-GLuc ON promotor/SEAP bi-cistronic vector. Four hours post-transfection, media was replaced and cells were maintained in either self-renewal media (wells plated with 10,000 cells) or differentiation media (wells plated with 20,000 cells). At indicated time, the secreted GLuc and SEAP activities were measured using the Secrete-Pair Dual Luminescence Assay Kit (GeneCopoeia), according to manufacturer’s protocol using a plate reader (SpectraMax i3, Molecular Devices).

### Yeast steady-state microscopy assay

Cells were grown overnight in raffinose synthetic media at 28 °C before dilution to an optimal density of 0.1–0.3 at 600 nm in galactose synthetic media. Cells were grown for 4-6 h at 28 °C to induce expression of the galactose-inducible Ubc9-2:eGFP, and subsequently shifted to 37 °C. After 30 min, an aliquot of the cells was centrifuged, washed once in water, and resuspended prior to mounting on glass slides. Fluorescence was visualized using a Zeiss Axiovert 200 inverted epifluorescence microscope and 100x/1.3 N.A. oil-immersion objective (Olympus), and a digital Axiocam MRm camera (Carl Zeiss) controlled with Axiovision software.

### Rate of protein synthesis

Protein synthesis was monitored by incorporation of O-propargyl-puromycin (OPP) (Blanco et al., 2016; Liu et al., 2012). NSPCs and differentiated progeny were labeled with 5 µM OPP for 30 min to label translating nascent chains. Controls for each sample were pretreated with cycloheximide (CHX; 100 µg/mL) for 30 min prior to OPP addition to correct for non-ribosomal background labeling or left unlabeled. Cells were washed twice in PBS, gently collected by dissociation with Accutase, and washed in PBS by centrifugation (200 g, 5 min, 4 °C). Cell pellets were resuspended and fixed in 1.6% PFA for 10 min at room temperature, washed, and then permeabilized with ice-cold methanol for 15 min at 4 °C. Samples were labeled by incubation with a click-chemistry reaction mix comprised of 5 µM Cy5-picolyl-azide, 0.5 mM Cu/BTTAA complex, 5 mM aminoguanidine, 100 mM sodium ascorbate in PBS for 30 min. After labeling, cells were washed into FACS Buffer (PBS, 1% BSA) and fluorescence was measured by flow cytometry.

## QUANTIFICATION AND STATISTICAL ANALYSIS

### Cell density

Cell density was measured in a Countess II (Life Technology).

### Viability Assays

For MTT viability assays, formazan absorbance measured for each sample was corrected for the total cell number/well (Hoechst fluorescence). Final viability scores were normalized to basal conditions. Analysis of 7-AAD FACS data was performed using FlowJo and Excel software.

### RNA-Seq analysis

Reads were mapped using Kallisto (Bray et al., 2016), and analyzed in R using the Sleuth package. Gene-level p-values for differential expression were calculated using the Wald test.

### REACTOME Enrichment Analysis

We estimated p-values by comparing the average change in expression in each pathway from our RNA-Seq analysis to the mean change of random chosen sets of genes of the same size. Estimates are based on up to 10^^5^ samples per term. The resampling was implemented in R using the script, DogGO.R (github.com/ptdolan/dogGO).

### GO Biological Processes Enrichment Analysis

Enrichment of specific biological processes in NPSCs and Differentiated Cells potentially altered ubiquitin-related genes (based on Uniprot annotations). Enrichment of specific GO Biological Processes (BPs) were assessed using the Panther database using Fisher’s Exact Test to assess over enrichment. Significance was determined based on a FDR corrected p-value (Mi et al., 2013).

### RT-qPCR

The relative expression of each sample was measured by the CFX software using the comparative 2ΔΔC_t_ method, and normalized to the mean expression of the housekeeping genes *Gusb* and *Gapdh* from replicates. Each probe was analyzed in duplicates for at least three independent samples per group.

### Western Blot Analysis

Blots were scanned using the LI-COR laser-based image detection method (Odyssey CLx scanner), and analyze using the Image Studio Lite Software (LI-COR Biosciences). For chaperone expression experiments (Figures 3E; 4A, B), same samples were used and results were thus verified in the same sample.

### Proteostat analysis

Microscopic monitoring of endogenous amyloid-like aggregates by Proteostat (Enzo Life Sciences) was done according to manufacturer’s instructions. For quantification of ICC Proteostat data, three coverslips were stained per condition per experiment, and at least 40 cells per coverslip were counted. Post-imaging analyses were done using ImageJ software. Imaged channels were split, maximal projections of confocal z-stacks taken (0.45 µm z-steps for 3 µm sections on average) were made followed by local background subtraction (rolling ball radius; same settings were applied to all experimental conditions), and merged. For quantification of fluorescence per cell, threshold was set to outline the positive areas followed by image conversion to black and white using the binary function. This value was divided by the sum of nuclei imaged for each sample.

Proteostat IHC imaging data analysis was done by importing raw image stacks into Fiji/ImageJ (NIH) software. The intensity value was measured, and normalized to background levels of each channel using unstained sections from aged-matched animals (rolling ball radius; same settings were applied to all experimental conditions).

Proteostat raw signal fluorescence intensities measured by a fluorimeter were corrected for the number of cells (Hoechst fluorescence).

### Yeast steady-state Microscopy Analysis

Quantification of yeast complementation assay represents three independent experiments, in which biological triplicates constitute different transformations. Image analyses was performed by ImageJ software. Only defined bright puncta were counted in at least 70 cells per experiment.

### Confocal Image Analysis

Unless stated otherwise, microscopic images (16-bit) displayed were acquired using an LSM700 or LSM880 AxioObserver confocal microscope (Carl Zeiss, Jena, Germany) by sequential excitation at 647, 568, 488 and 408 nm. The same exposure settings were used across all conditions in each individual experiment. Acquisition parameters included a speed of 400 Hz, pixel dwell time of 600 ns, line averaging of 2. Tile scans and z-stacks of the area of interest were obtained and stitched using the ZEN Black software. Fiji (ImageJ) was used for image processing. Channels were split and image backgrounds were subtracted (rolling ball radius between 50 – 150 pixels), and channels were subsequently merged. The same settings were applied to all experimental conditions. For signal quantification, thresholds were set to outline the positive areas and images were converted to black and white using the binary function; gaps were filled using the watershed function. The same threshold values were used for all images across all conditions in each individual experiment. Z-stack projection were generated in Fiji using the same settings for all images for each experiment. For visualization of some images displayed, brightness was adjusted (post-quantification analyses) in Fiji to enhance visualization. The same settings were applied to all images shown for each experiment. Unless stated otherwise, scale bars represent 10 μm

### Luciferase-promoter based activity analysis

Raw values of the promotor activities were normalized and calculated as ratio of GLuc to SEAP activities.

### Statistical analysis

Unless state otherwise in the above, all quantitative results presented represent mean ± SD of at least three independent experiments performed, in which biological triplicates constitute three different NSPC isolations. In each experiment technical replicates were included. No statistical methods were used to predetermine sample size. Statistical comparisons were made by (two-tailed) unpaired t-test assuming Gaussian distribution with Welch’s correction and one-way analysis of variance (ANOVA), as appropriate, using the GraphPad Prism 7.0 Software. Differences were considered to be significant at *p* < 0.05. * *p* < 0.05; ** *p* < 0.01; *** *p* < 0.005; **** *p* < 0.001, unless stated otherwise.

## DATA AND CODE AVAILABILITY

All data to support the conclusions in this manuscript can be found in the figures, and are available from the corresponding authors upon reasonable request. RNA-Seq reads have been deposited to the NCBI short read archive under Bioproject PRJNA531489 (http://www.ncbi.nlm.nih.gov/bioproject/531489). Associated data tables can be found at the Stanford Data Repository (https://purl.stanford.edu/gd935jm2479). The resampling of the REACTOME enrichment analysis was implemented in R using the script, DogGO.R (github.com/ptdolan/dogGO). Original imaging data files have been deposited in Mendeley Data (http://dx.doi.org/10.17632/sgdg7bv2dw.1).

## Supplemental item titles

**Table S2. Differential gene expression between NSPCs and differentiated neural progeny cells** (Excel document (.csv); *Related to Figures 3, S3*)

**Table S3. REACTOME enriched terms in NSPCs *vs*. differentiated neural progeny cells** (Excel document (.csv); *Related to Figures 3, S3*)

