## Supplemental Information (Figures S1-S7; supplemental tables, Data S1) for "Differentiation drives widespread rewiring of the neural stem cell chaperone network"

This file includes:

- Figures S1-S7
- Legends to Supplemental Figures
- Tables S1, S4
- Data S1

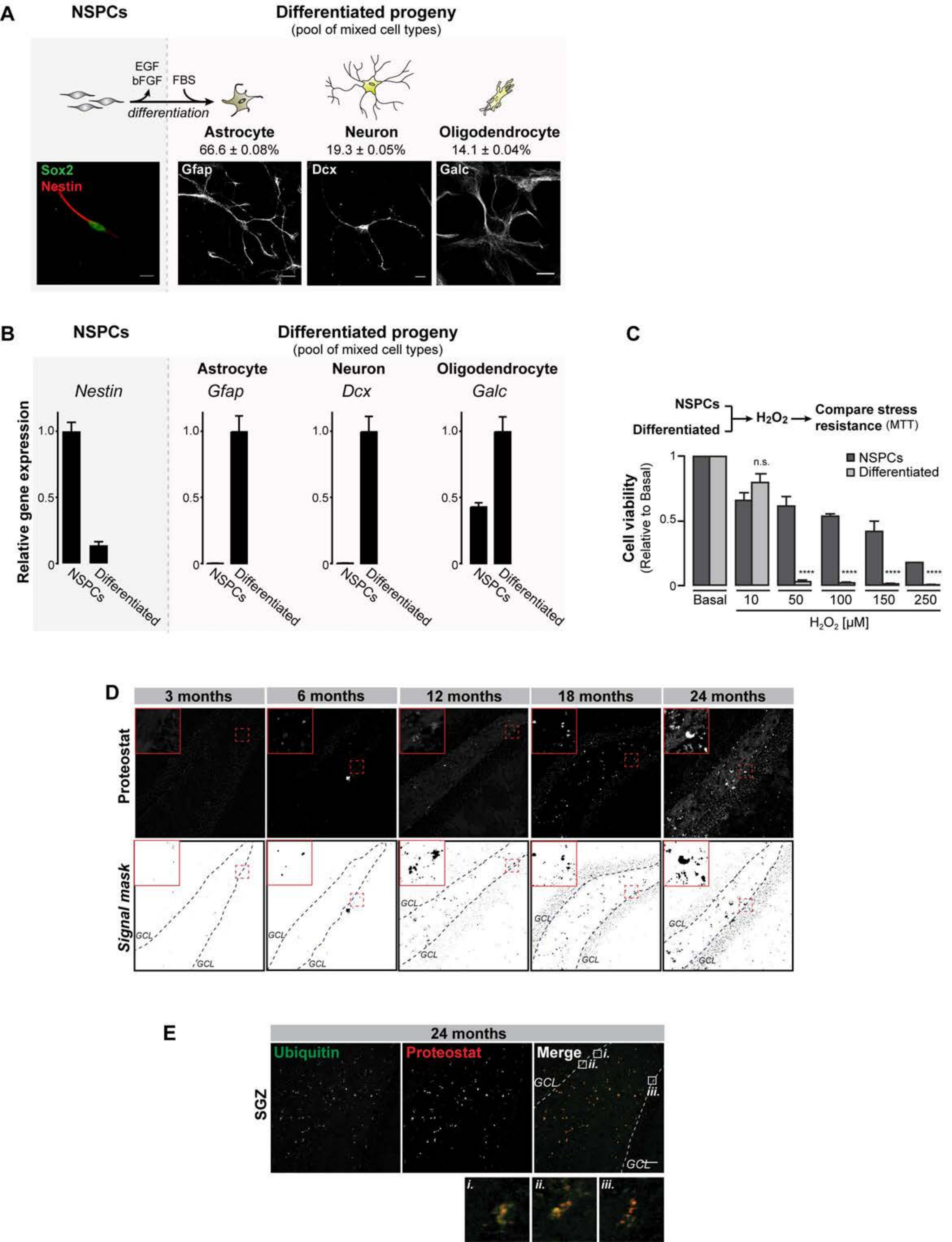
A
NSPCs
Differentiated progeny
(pool of mixed cell types)
EGF
bFGF
FBS
differentiation
Astrocyte
66.6 ± 0.08%
Neuron
19.3 ± 0.05%
Oligodendrocyte
14.1 ± 0.04%
Sox2
Nestin
Gfap
Dcx
Galc
B
NSPCs
Differentiated progeny
(pool of mixed cell types)
Astrocyte
Neuron
Oligodendrocyte
Nestin
Gfap
Dcx
Galc
Relative gene expression
NSPCs
Differentiated
C
NSPCs
Differentiated
H2O2
Compare stress resistance (MTT)
Cell viability
(Relative to Basal)
NSPCs
Differentiated
n.s.
****
Basal
10
50
100
150
250
H2O2 [μM]
D
3 months
6 months
12 months
18 months
24 months
Proteostat
Signal mask
GCL
E
24 months
Ubiquitin
Proteostat
Merge
SGZ
GCL
i.
ii.
iii.

Figure S2 - Vonk *et al.*

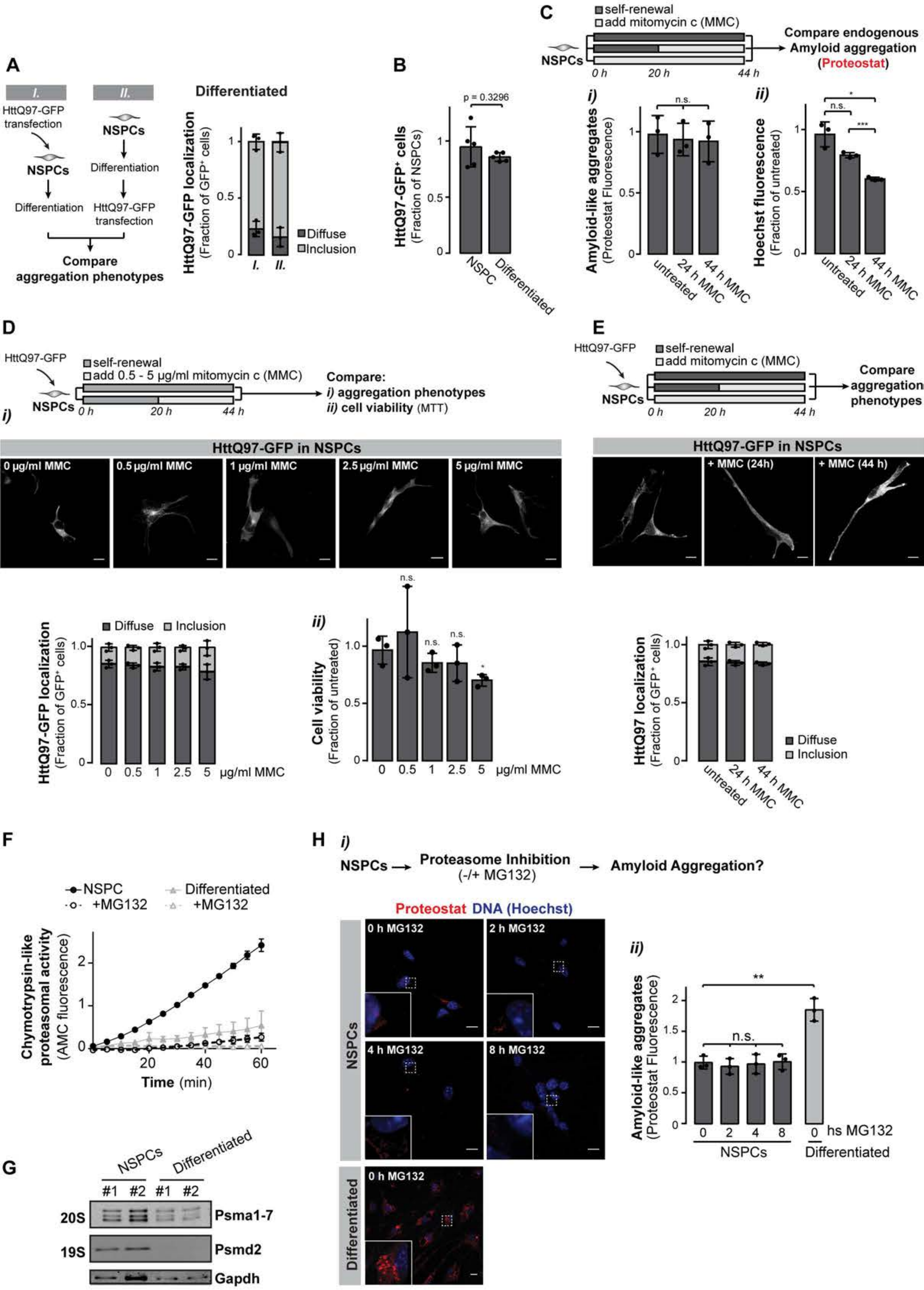

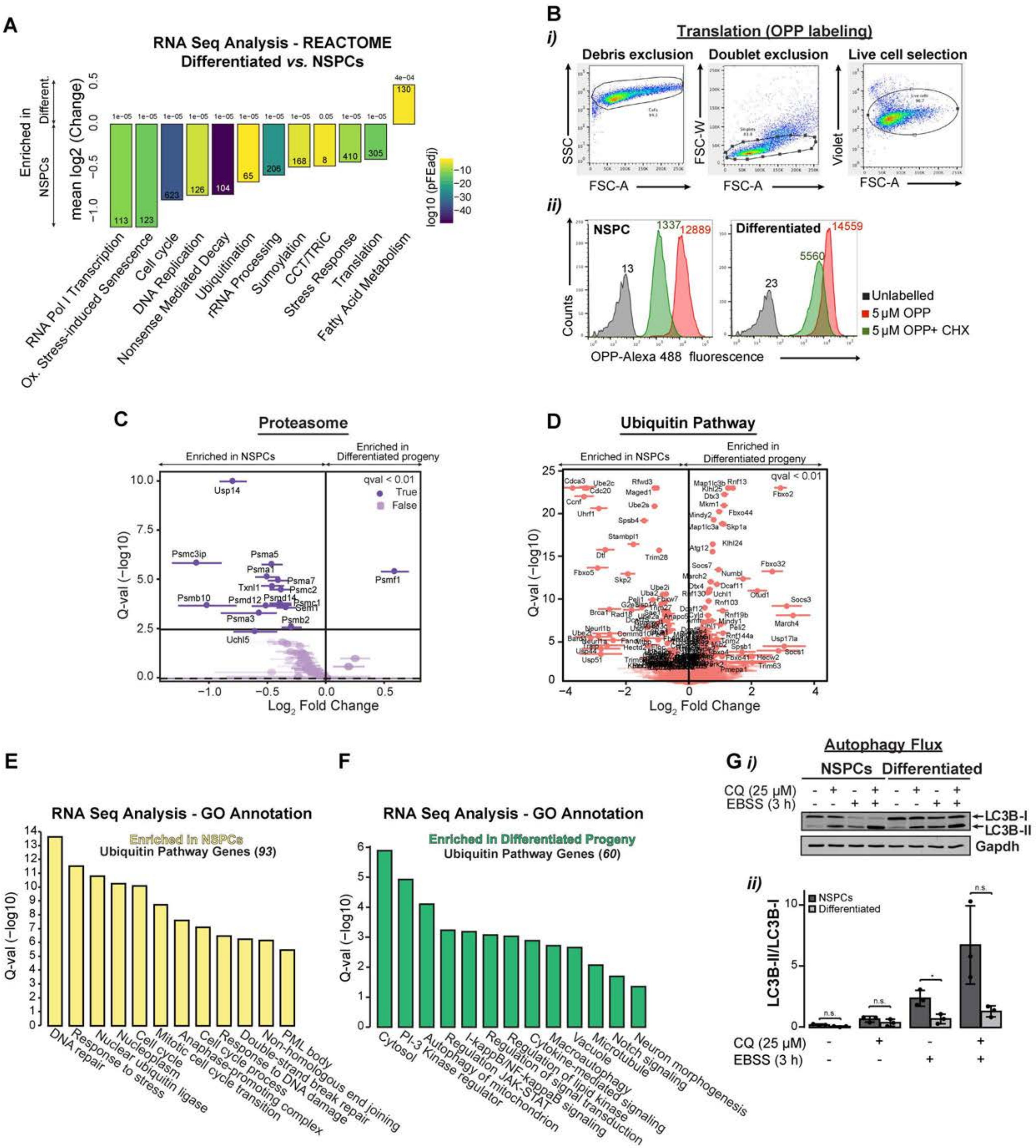
A
RNA Seq Analysis - REACTOME
Differentiated vs. NSPCs
Enriched in NSPCs
Different.
mean log2 (Change)
log10 (pFEadj)
RNA Pol I Transcription
Ox. Stress-induced Senescence
Cell cycle
DNA Replication
Nonsense Mediated Decay
Ubiquitination
rRNA Processing
Sumoylation
CCT/TRiC
Stress Response
Translation
Fatty Acid Metabolism
B
i)
Translation (OPP labeling)
Debris exclusion
Doublet exclusion
Live cell selection
SSC
FSC-W
Violet
FSC-A
ii)
NSPC
Differentiated
Counts
OPP-Alexa 488 fluorescence
Unlabelled
5 µM OPP
5 µM OPP+ CHX
C
Proteasome
Enriched in NSPCs
Enriched in Differentiated progeny
qval < 0.01
True
False
Q-val (−log10)
Log2 Fold Change
D
Ubiquitin Pathway
E
RNA Seq Analysis - GO Annotation
Enriched in NSPCs
Ubiquitin Pathway Genes (93)
DNA repair
Response to stress
Nuclear ubiquitin ligase
Nucleoplasm
Cell cycle
Mitotic cell cycle transition
Anaphase-promoting complex
Cell cycle process
Response to DNA damage
Double-strand break repair
Non-homologous end joining
PML body
F
RNA Seq Analysis - GO Annotation
Enriched in Differentiated Progeny
Ubiquitin Pathway Genes (60)
Cytosol
PI-3 Kinase regulator
Autophagy of mitochondrion
Regulation JAK-STAT
I-kappaB/NF-kappaB signaling
Regulation of signal transduction
Regulation of lipid kinase
Cytokine-mediated signaling
Macroautophagy
Vacuole
Microtubule
Notch signaling
Neuron morphogenesis
G i)
Autophagy Flux
NSPCs
Differentiated
CQ (25 µM)
EBSS (3 h)
LC3B-I
LC3B-II
Gapdh
ii)
NSPCs
Differentiated
LC3B-II/LC3B-I

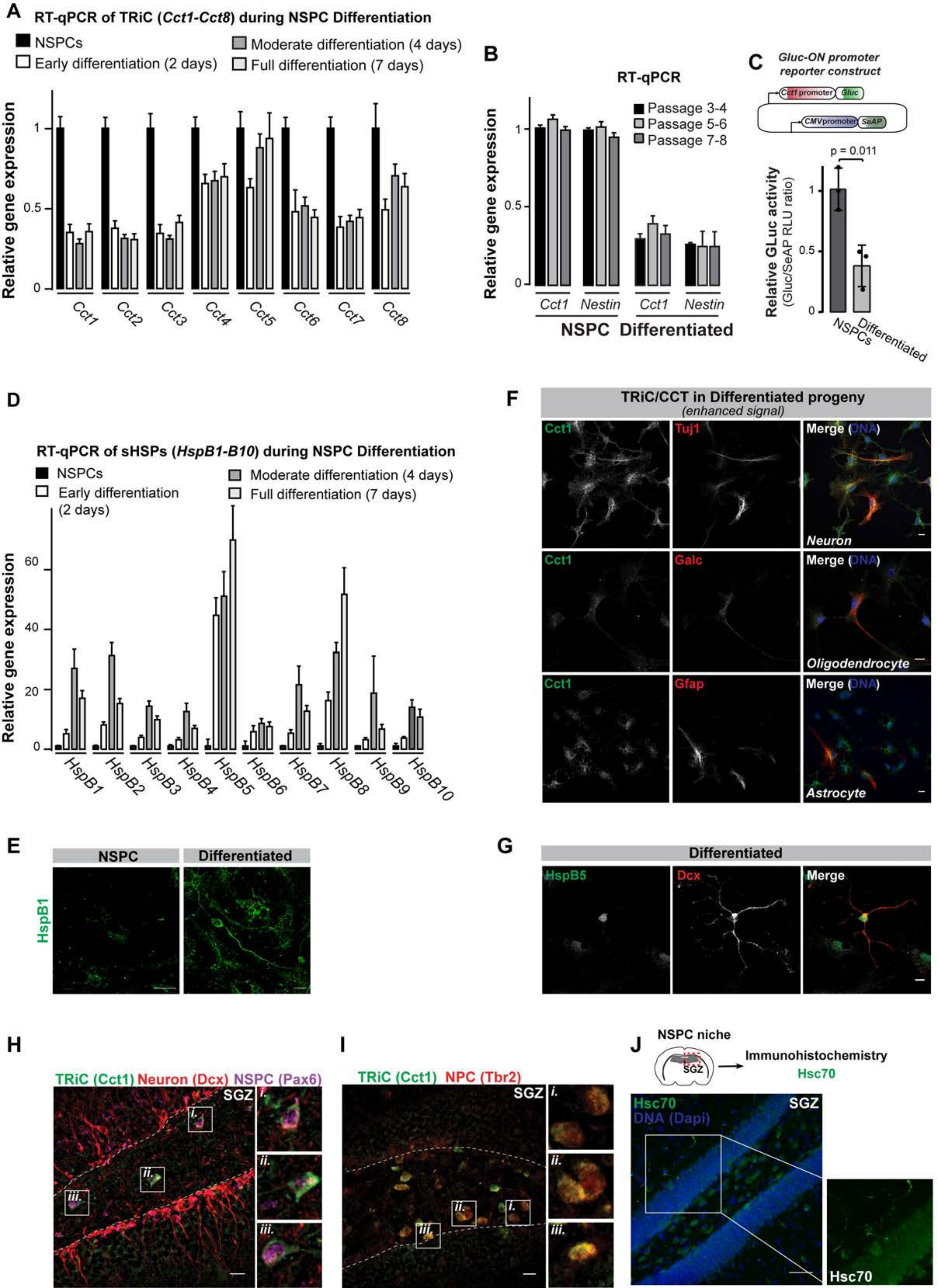
A
RT-qPCR of TRiC (Cct1-Cct8) during NSPC Differentiation
NSPCs
Early differentiation (2 days)
Moderate differentiation (4 days)
Full differentiation (7 days)
Relative gene expression
Cct1
Cct2
Cct3
Cct4
Cct5
Cct6
Cct7
Cct8
B
RT-qPCR
Passage 3-4
Passage 5-6
Passage 7-8
Cct1
Nestin
NSPC
Differentiated
C
Gluc-ON promoter reporter construct
Cct1 promoter
Gluc
CMV promoter
SeAP
p = 0.011
Relative GLuc activity (Gluc/SeAP RLU ratio)
NSPCs
Differentiated
D
RT-qPCR of sHSPs (HspB1-B10) during NSPC Differentiation
NSPCs
Early differentiation (2 days)
Moderate differentiation (4 days)
Full differentiation (7 days)
HspB1
HspB2
HspB3
HspB4
HspB5
HspB6
HspB7
HspB8
HspB9
HspB10
E
NSPC
Differentiated
HspB1
F
TRiC/CCT in Differentiated progeny
(enhanced signal)
Cct1
Tuj1
Merge (DNA)
Neuron
Galc
Oligodendrocyte
Gfap
Astrocyte
G
Differentiated
HspB5
Dcx
Merge
H
TRiC (Cct1) Neuron (Dcx) NSPC (Pax6)
SGZ
I
TRiC (Cct1) NPC (Tbr2)
J
NSPC niche
SGZ
Immunohistochemistry
Hsc70
DNA (Dapi)

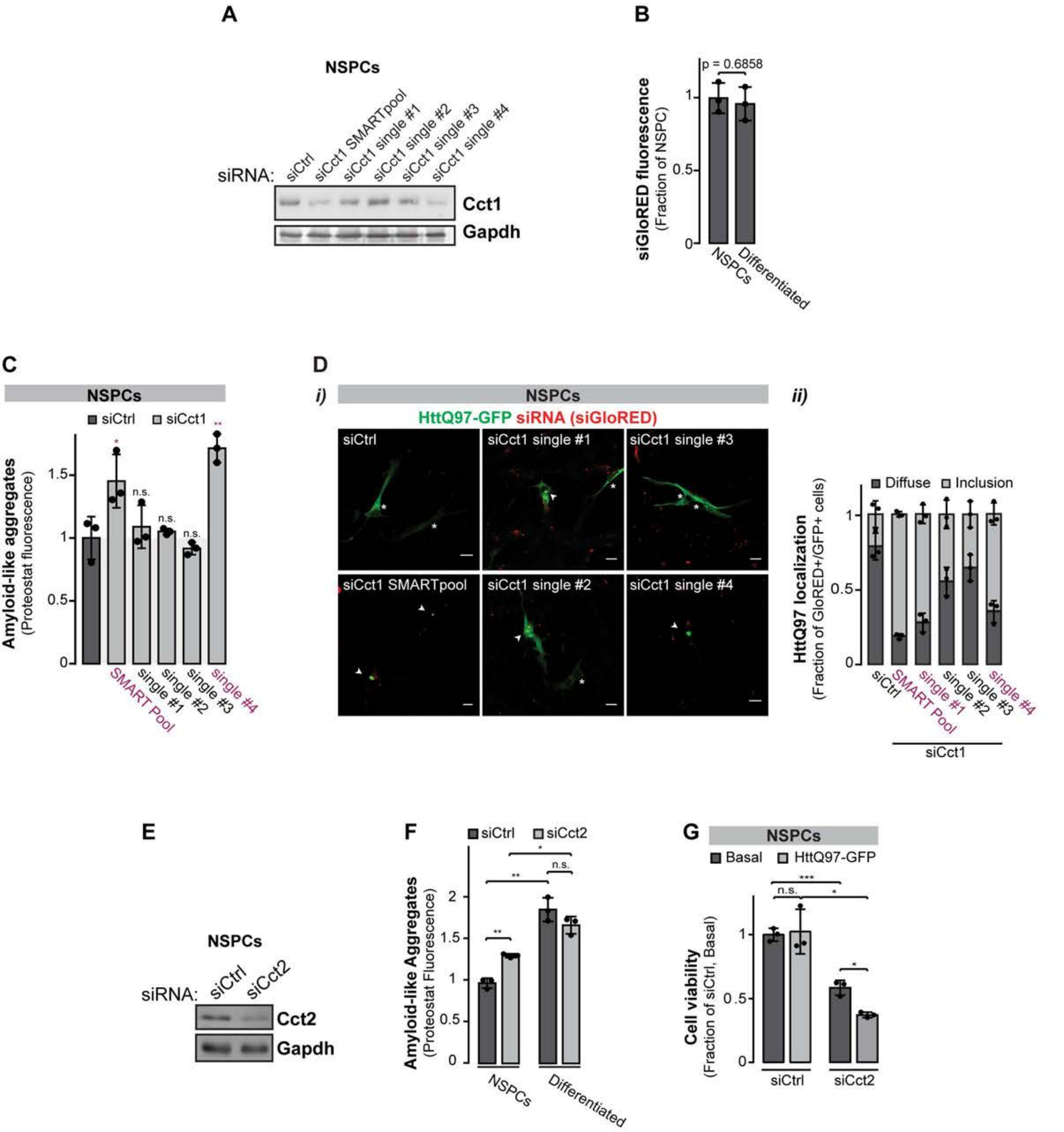
A
NSPCs
siRNA:
siCtrl
siCct1 SMARTpool
siCct1 single #1
siCct1 single #2
siCct1 single #3
siCct1 single #4
Cct1
Gapdh
B
p = 0.6858
siGloRED fluorescence
(Fraction of NSPC)
NSPCs
Differentiated
C
NSPCs
siCtrl
siCct1
Amyloid-like aggregates
(Proteostat fluorescence)
n.s.
SMART Pool
single #1
single #2
single #3
single #4
D
i)
NSPCs
HttQ97-GFP siRNA (siGloRED)
siCtrl
siCct1 single #1
siCct1 single #3
siCct1 SMARTpool
siCct1 single #2
siCct1 single #4
ii)
Diffuse
Inclusion
HttQ97 localization
(Fraction of GloRED+/GFP+ cells)
siCtrl
SMART Pool
single #1
single #2
single #3
single #4
siCct1
E
NSPCs
siRNA:
siCtrl
siCct2
Cct2
Gapdh
F
siCtrl
siCct2
n.s.
Amyloid-like Aggregates
(Proteostat Fluorescence)
NSPCs
Differentiated
G
NSPCs
Basal
HttQ97-GFP
n.s.
Cell viability
(Fraction of siCtrl, Basal)
siCtrl
siCct2

Figure S6 - Vonk *et al*.

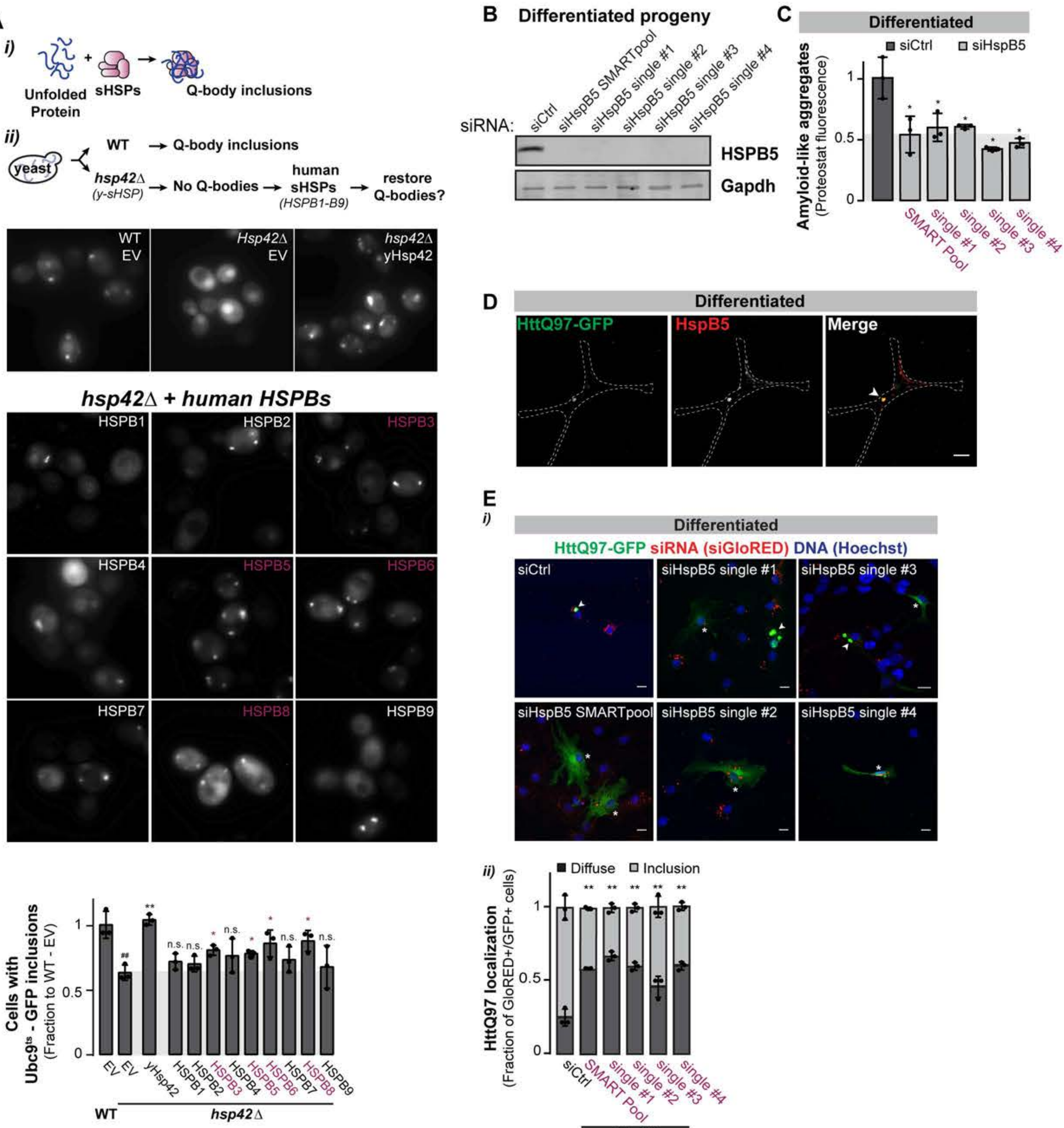

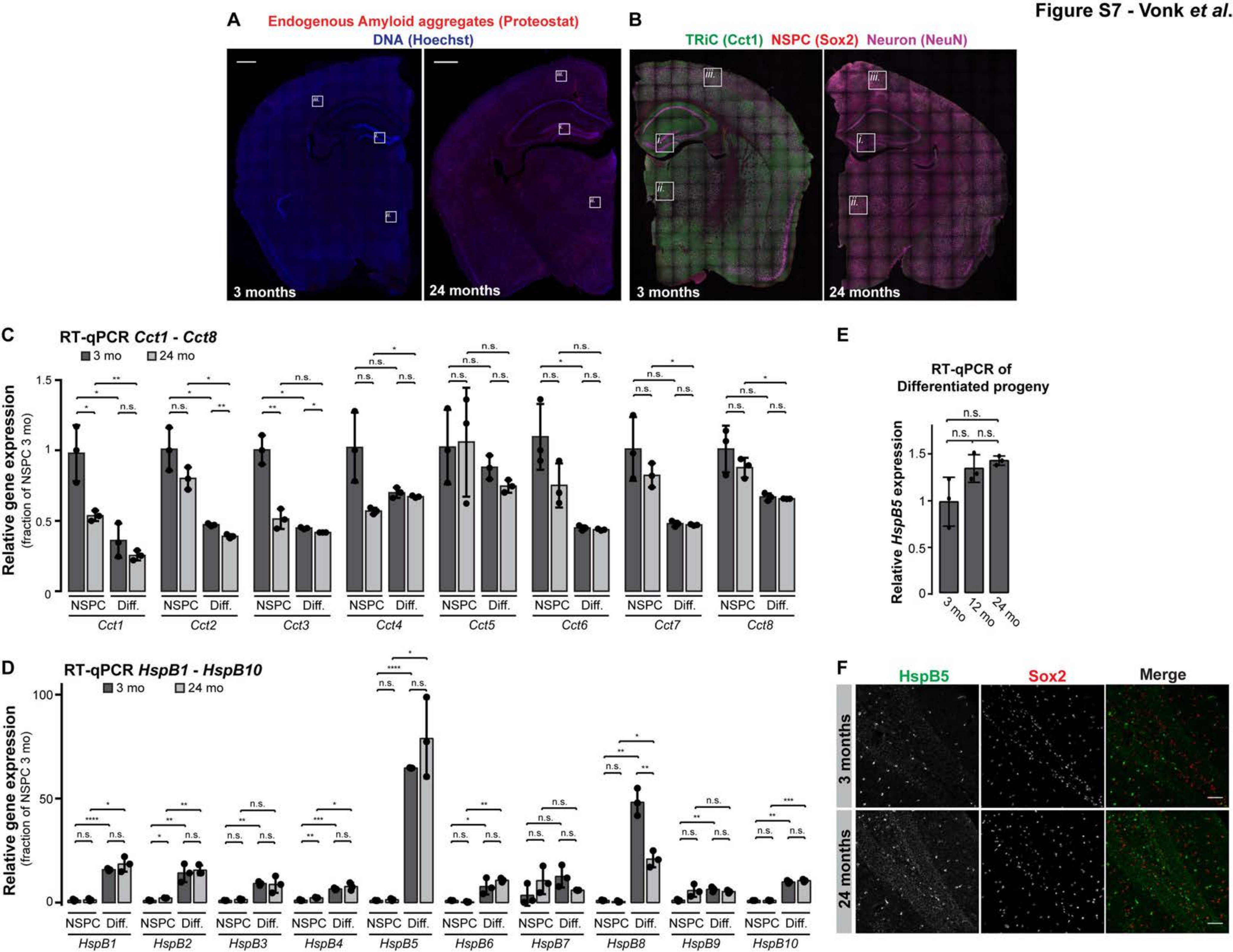

Figure S7 - Vonk et al.
A
Endogenous Amyloid aggregates (Proteostat)
DNA (Hoechst)
3 months
24 months
B
TRiC (Cct1) NSPC (Sox2) Neuron (NeuN)
3 months
24 months
C
RT-qPCR Cct1 - Cct8
3 mo
24 mo
Relative gene expression
(fraction of NSPC 3 mo)
NSPC
Diff.
Cct1
Cct2
Cct3
Cct4
Cct5
Cct6
Cct7
Cct8
D
RT-qPCR HspB1 - HspB10
3 mo
24 mo
Relative gene expression
(fraction of NSPC 3 mo)
HspB1
HspB2
HspB3
HspB4
HspB5
HspB6
HspB7
HspB8
HspB9
HspB10
E
RT-qPCR of
Differentiated progeny
Relative HspB5 expression
3 mo
12 mo
24 mo
F
HspB5
Sox2
Merge
3 months
24 months

### Legends to Supplemental Figures

**Figure S1. Brain aging is associated with an increase in amyloid-like aggregates in the SGZ NSPC niche** *(Related to Figure 1)*

**A,** Sox2$^+$/Nestin$^+$ NSPCs, obtained via crude isolation from SGZ and SVZ NSC niches in the mouse brain, were cultured under self-renewal conditions (i.e. in presence of the growth factors EGF and bFGF). Differentiation was induced by growth factor removal, plain media-wash, and addition of 0.5 % FBS to the culture media as described in Methods, and resulted in a mixed pool of differentiated cells (in which three main neural lineages were represented as follows (mean ± SD): ~ 67 % astrocytes (Gfap$^+$), ~ 19 % neurons (Dcx$^+$), and ~ 14 % oligodendrocytes (Galc$^+$)). Confocal imaging for NSPC and the three main differentiated neural lineages upon immunostaining with indicated antibodies. Scale bars: 10 µm. For each lineage, positive cells were manually counted. Data was collected for three independent experiments of at least 80 cells each.

**B,** mRNA expression of cellular markers in NSPCs and differentiated progeny (NSPC: *Nestin*; Astrocyte: *Gfap*; Neuron: *Dcx*; Oligodendrocyte: *Galc*), analyzed in NSPCs and the pool of differentiated cells containing mixed cell types by RT-qPCR. The data represent mean ± SD of three independent experiments with quadruplicate measurements for each condition. Plotted values were scaled to the range [0,1] for visualization of all genes on the same axis.

**C,** Brain aging is associated with increased amyloid-like aggregation in the hippocampus. Confocal imaging of Proteostat-reactive deposits in sections of the SGZ region in the hippocampus of brains of 3, 6, 12, 18, and 24 month old mice. Top: Proteostat fluorescence (white); bottom: Signal masks of Proteostat fluorescence. Dotted lines illustrate borders of SGZ/GCL. Scale bars: 50 µm. Insets: Magnification of Proteostat-reactive amyloid-like aggregate deposition in SGZ of hippocampus (red square).

**D,** Confocal imaging of ubiquitin$^+$ (green)/Proteostat$^+$ (red) inclusions in the dentate gyrus of the hippocampus of the aged brain (24 month old). Single and merge images are shown. Dotted lines illustrate borders of SGZ/GCL. Scale bar: 50 µm. Insets: Magnification of the ubiquitin$^+$/Proteostat$^+$ deposits in the SGZ in the aged brain.

**Figure S2. Distinct proliferation rates of NSPCs and differentiated cells do not affect cellular ability to maintain proteome solubility** *(Related to Figure 2)*

**A,** HttQ97-GFP transfection prior or after NSPC differentiation does not alter the cellular localization of misfolded disease-linked HttQ97-GFP in differentiated neural cells. HttQ97-GFP cellular localization was studied under two conditions: (***I.***) HttQ97-GFP was transfected into NSPCs. Four hours post-transfection, media was replaced and NSPC differentiation was induced; (***II.***) NSPC were differentiated for four days prior to transfection with HttQ97-GFP. Media was replaced 4 hours post-transfection. In both conditions, cells were fixed after 7 days of differentiation. Graph (mean ± SD) shows quantification of diffusely localized versus aggregated HttQ97-GFP in differentiated neural cells, based on confocal imaging. Data was collected from three independent experiments of at least 40 cells each.

**B,** HttQ97-GFP is equally expressed in NSPCs and differentiated progeny. Relative fluorescence of HttQ97-GFP in NSPCs and differentiated progeny cells upon transient transfection of HttQ97-GFP in NPSCs and cell maintenance under self-renewal (NSPCs) or differentiation (differentiated) conditions, measured by Flow cytometry as $GFP^+$ cells (10,000 total events per sample were counted). Data are graphed as mean ± SD. $p = 0.3296$ (non-significant).

**C,** Impaired rate of NSPC proliferation by mitomycin C (MMC) treatment does not change the abundance of endogenous amyloid-like aggregates in NSPCs. NSPCs were treated with 1 µg/ml MMC for indicated time points prior to Hoechst and Proteostat stainings, and fluorogenic measurements. (***i***) Graph shows quantification of Proteostat fluorescence, normalized to untreated conditions and corrected for the number of cells (Hoechst stain). (***ii***) MMC was used at an effective concentration as treatment resulted in a decline in the total cell number, determined by nuclei count (Hoechst fluorescence). Data were collected from three independent experiments; bars show mean ± SD. Value of untreated was arbitrarily set at 1. * $p < 0.05$; *** $p < 0.005$; n.s., non-significant.

**D,** Inhibition of NSPC proliferation by mitomycin C (MMC) does not result in HttQ97-GFP aggregate formation nor reduced cell viability. (***i***) Confocal imaging of HttQ97-GFP in NSPC treated for 24 hours with increasing concentrations of MMC (0 – 5 µg/ml). Scale bars: 10 µm. Graph shows quantification of diffusely localized versus aggregated protein in NSPCs at distinct MMC concentrations. Data was collected from three independent experiments of at least 40 cells per condition each; bars show mean ± SD. (***ii***) Comparison of metabolic activity of viable NSPCs upon treatment with increasing concentrations MMC. NSPCs were treated for 24 hours with similar concentrations of MMC as in (***i***). Cell viability (i.e. formazan absorbance) is shown as mean ± SD of three independent experiments with four replicates per dose. Total cell number per well was calculated prior to MTT addition by nuclei measurement upon Hoechst fluorogenic staining; MTT absorbance for each sample was corrected for the total cell number/well. Value of untreated NSPCs was arbitrarily set at 1. * $p < 0.05$; n.s., non-significant.

**E,** Attenuation of NSPC proliferation rate by mitomycin C (MMC) treatment does not change abundance of HttQ97 aggregates. Confocal imaging of HttQ97-GFP in NSPCs treated with 1 µg/ml MMC for 0, 24, and 44 h prior to fixation. Scale bars: 10 µm. Graph shows quantification of diffusely localized versus aggregated HttQ97-GFP in untreated and MMC-treated NSPCs. Data was collected from three independent experiments of at least 60 cells per condition each; bars show mean ± SD.

**F,** Chymotrypsin-like peptidase activity of the 20S proteasome core is significantly higher in NSPCs than in differentiated progeny. Graph shows proteasomal activity for NSPCs and differentiated progeny at basal (solid lines) and proteasome-inhibited (+ 10 µM MG132; dashed lines) conditions, determined by quantitative measurement of fluorescent AMC release from the N-Suc-Leu-Leu-Val-Tyr-AMC peptide substrate upon proteasomal cleavage. Quantified data were collected from three independent experiments and represented as mean ± SD.

**G,** NSPC differentiation induces a decreased expression of Psmd2, the 19S subunit of the 26S proteasome responsible for poly-ubiquitylated substrate recognition. Immunoblot analysis with antibodies for Psma1-7 (20S catalytic core subunit), Psmd2 (19S regulatory subunit), and Gapdh (loading control) in NSPCs and differentiated progeny. Biological duplicates were loaded.

**H,** Blockage of proteasome in NSPCs does not lead to increased abundance of endogenous aggregates. (***i***) Confocal imaging of amyloid-like aggregate formation in NSPCs and differentiated progeny at basal (0 h) or at proteasome-inhibited conditions (+ MG132, 15 µM for indicated hours prior to fixation). Scale bars: 10 µm. Insets: Magnification of Proteostat$^{+}$-deposits (square). (***ii***) Graph shows quantified Proteostat fluorescence (Ex: 550 nm, Em: 600 nm) in NSPCs and differentiated progeny, corrected for the amount of cells (Hoechst fluorescence). Data was collected from three independent experiment, and graphed as mean ± SD. ** $p < 0.01$; n.s., non-significant.

**Figure S3. Differential activation of ubiquitin-proteasome system during NSPC differentiation does not define cellular capacity to maintain proteome solubility**

*(Related to Figure 3)*

**A,** Bar graph showing the log2 mean effect of differentiation on gene expression in key enriched pathways (based on over-representation - Benjamini-Yekutieli-Corrected Fisher's Exact Test p<0.05) from the REACTOME databases. Bars are labeled with the number of genes in the pathway and significance of the mean change, compared to 100,000 randomly sampled gene sets of the same size.

**B,** Overall bulk rate of protein synthesis is reduced in differentiated neural progeny. (***i***) Gating for measurement of protein synthesis rates by Flow cytometry. Using the forward scatter (FSC) and side scatter (SSC) counts, cell debris and doublets were excluded from the measurement. The single, viable cells were gated using Violet Live/Dead Cell stain. (***ii***) Bulk rates of protein synthesis were determined in NSPCs and differentiated neural cells by incorporation of O-propargyl-puromycin (OPP) into newly translated proteins (Blanco et al., 2016; Liu et al., 2012). Flow cytometric plots of OPP fluorescence per counts at basal or protein synthesis inhibited (100 µg/mL cyclohexamide (CHX), 30 min prior to OPP addition) conditions. For quantification of relative rates of protein synthesis, see Figure 3C.

**C,** Proteasomal transcripts are underrepresented in the differentiated neural cell population. Volcano plot of differentially expressed proteasome genes in differentiated progeny cells compared to NSPCs. X-axis values are $\log_2$ (fold change), y-axis values are the –log10 of q-value. Q-values indicate FDR-adjusted p-value for multiple testing. Each point represent the indicated gene.

**D,** Ubiquitin pathway transcripts are heterogeneously expressed in NSPCs and their differentiated progeny. Volcano plot of differentially expressed genes involved in various ubiquitin pathways in differentiated progeny cells compared to NSPCs. X-axis values are $\log_2$ (fold change), y-axis values are the –log10 of q-value. Q-values indicate FDR-adjusted p-value for multiple testing. Each point represent the indicated gene.

**E, F,** Bar plots of GO ubiquitin pathway transcripts, enriched in either NSPCs (**E**) or their differentiated progeny (**F**). Mean of each category is plotted as -log10 of q-value.

**G,** Autophagy flux is reduced in differentiated neural progeny compared to NSPCs. (***i***) Autophagy flux is detected by LC3-II turnover using immunoblot analysis at basal conditions, in the presence of the lysosomal degradation inhibitor chloroquine (CQ, 25 μM for 3 h prior to lysis), or under starvation conditions (EBSS, 3 h prior to lysis). Upon increasing autophagic flux, the level of LC3-II will be increased in the presence of CQ, EBSS, or the combined treatment. Gapdh serves as loading control as indicated. (***ii***) Graph shows quantified LC3-II turnover in NPSCs and differentiated progeny at indicated conditions. Data was collected from three independent experiments, and graphed as mean ± SD. Value of untreated NSPCs was arbitrarily set at 1. * $p < 0.05$; n.s., non-significant.

**Figure S4. NSPC differentiation remodels the cellular chaperone network from high TRiC/low sHSP expression in NSPCs to low TRiC/high sHSP levels in differentiated neural progeny** *(Related to Figures 3 and 4)*

**A,** mRNA expression of *Cct1*-*Cct8* during NSPC differentiation (early (day 2); moderately (day 4); fully (day 7) maturated), analyzed by RT-qPCR. The data represent mean ± SD of three independent experiments with quadruplicate measurements for each condition, corrected for the expression of the housekeeping genes *Gusb* and *Gapdh*. For each gene studied, mRNA abundance in NSPCs was set at 1.

**B,** Loss of TRiC/CCT expression during NSPC differentiation is not an effect of artificial cell culture conditions influenced by passage number. mRNA expression of *Cct1* and that of the NSPC marker *Nestin* do not decline with increasing NSPC passage number, determined by RT-qPCR in NSPCs and 7 day differentiated neural cells and analyzed as described under **A**.

**C,** *Cct1* (GLuc) and CMV (SEAP; secreted tracking gene for data normalization) promoter activities were determined in NSPCs and differentiated neural progeny cell supernatants by measurement of their relative light units (RLU) following expression of the GLuc-ON *Cct1* promoter reporter construct in NSPCs prior to differentiation. Graph represent relative GLuc activities (mean ± SD) of three independent experiments with quadruplicate measurements for each condition; *Cct1* promoter activity of differentiated neural progeny cells was normalized to NSPC values and calculated as ratio of GLuc to SEAP activities. $p = 0.011$.

**D,** mRNA expression of *HspB1 - HspB10* in NSPCs and during NSPC differentiation (early (day 2); moderately (day 4); fully (day 7) maturated), analyzed by RT-qPCR. The data represent mean ± SD of three independent experiments with quadruplicate measurements for each condition, corrected for the expression of the housekeeping genes *Gusb* and *Gapdh*. For each gene studied, mRNA abundance in NSPCs is set at 1.

**E,** Confocal images of HspB1 (green) in NSPCs and 7-day differentiated neural progeny cells. Scale bars: 10 µm.

**F,** Cct1 is similarly attenuated in all three main differentiated neural lineages following NSPC differentiation. Confocal imaging of Cct1 (green) in the distinct cell lineages of the differentiated population, identified by specific cell markers (red): Gfap for astrocytes, Tuj1 for neurons, and Galc for oligodendrocytes. DNA (Hoechst) in blue. For visualization purposes, the low Cct1 fluorescent signal in differentiated neural cells was enhanced during post-imaging analysis. No clear cellular differences in Cct1 expression were observed within the different cell types of the differentiated neural cell population. Scale bars: 10 µm.

**G,** HspB5 is expressed in $Dcx^{+}$ neuronal cells obtained by differentiation of mouse brain-isolated NSPCs. Co-immunostaining of HspB5 (green) with neuronal marker Dcx (red) followed by confocal imaging (single and merged images are shown). Scale bars: 10 µm.

**H, I,** Mouse brain sections of the SGZ NSPC niche, immunostained for (**G**) Dcx (red; indicates neurons), Pax6 (magenta, indicates NSPCs), and TRiC (Cct1, green), or (**H**) Tbr2 (red, indicates NSPCs) and TRiC (Cct1, green). Scale bars: 50 µm. Insets: Magnification of $Pax6^{+}/Cct1^{+}$ (**G**) or $Tbr2^{+}/Cct1^{+}$ (**H**) cells in the SGZ region at indicated areas (white squares); single and merge channels are displayed. Dotted lines illustrate borders of SGZ/GCL.

**J,** Hsc70 is heterogeneously expressed in the SGZ of the young adult brain. Confocal image of brain section of the SGZ immunostained for invariant Hsc70 (green). Nuclei in blue (Dapi). Scale bar: 50 µm. Inset: Magnification of Hsc70 distribution in SGZ (white square; single channel).

**Figure S5. Depletion of TRiC/CCT results in increased aggregate abundance and vulnerability to cellular damage under proteotoxic stress conditions** *(Related to Figure 5)*

**A,** Immunoblot analyses of Cct1 in control (siCtrl)- and siCct1-targeted NSPCs. Cct1 expression was depleted by treatment with either a mixed pool of multiple siRNAs (SMARTpool) or single siRNAs (#1 - #4). Gapdh serves as loading control. Degree of Cct1 knockdown is representative for all experiments presented. Note: not all single siRNAs were as efficient in depleting Cct1 from NSPCs as the siRNA SMARTpool.

**B,** Fluorescence intensities of the siGloRED siRNA-transfection indicator were comparable in NSPC and differentiated cell populations, thus, differential levels of transfected siRNAs or a potential siRNA-dilution in proliferative NSPCs can be excluded as a significant contributing factor to the observed aggregation phenotypes. Data from three independent experiments, measured by microplate reading, and graphed as mean ± SD. SiGloRED fluorescence of NSPCs is set at 1. $p = 0.0708$ (non-significant).

**C,** Similar as to NSPCs treated with a mixed pool of multiple siRNAs targeting Cct1 (siCct1 SMART Pool), Cct1 depletion from NSPCs by single siRNA-treatment (single siRNA #1 - #4, light gray bars) resulted in increased aggregation of endogenous amyloid-like proteins; quantified by fluorescence (mean ± SD), corrected for the number of cells. Proteostat fluorescence of siCtrl-targeted NSPCs is set at 1. Significant inductions in Proteostat fluorescence upon Cct1 depletion are indicated in purple. * $p < 0.05$; ** $p < 0.01$; n.s., non-significant.

**D,** (***i***) Confocal imaging of HttQ97-GFP (green) and siGloRed (red; indicator of siRNA transfection) in control (siCtrl) and siCct1-treated NSPCs and differentiated neural cells. Cct1 expression was depleted by treatment with either a mixed pool of multiple siRNAs (SMART Pool) or single siRNAs (#1 - #4). Scale bars: 10 µm; arrows indicate aggregates, asterisks indicate diffusely expressed HttQ97-GFP. (***ii***) Quantification (mean ± SD) of diffusely localized versus aggregated HttQ97-GFP in double-transfected siCtrl/siGloRed$^{+}$- and siCct1/siGloRed$^{+}$-treated (SMARTpool *vs.* single siRNAs) NSPCs from at least 40 cells per condition per experiment. Significant inductions in the abundance of HttQ97-GFP inclusions upon Cct1 depletion are indicated in purple ($p < 0.05$).

**E,** Immunoblot analyses of Cct2 in control (siCtrl)- and siCct2-targeted NSPCs. Gapdh serves as loading control. Degree of Cct2 knockdown is representative for all experiments presented.

**F,** Aggregation of endogenous proteins in NSPC treated with control or a pool of siRNAs targeting Cct2; quantified by fluorescence (mean ± SD), corrected for the number of cells. Proteostat fluorescence of siCtrl-targeted NSPCs is set at 1. ** $p < 0.01$; **** $p < 0.001$; n.s., non-significant.

**G,** Decreased viability of HttQ97-GFP expressing NSPCs is further exacerbated by depletion of Cct2, similar as determined for Cct1 (Figure 5G). Cell viability in control (siCtrl) and siCct2-treated wild-type and HttQ97-GFP expressing cells was measured by MTT-based assays. Formazan absorbance for each sample was corrected for the total cell number/well prior to MTT addition by Hoechst staining of nuclei. Data from three independent experiments is graphed as mean ± SD. Viability of control cells under basal conditions is set at 1. n.s., non-significant; * $p < 0.05$; *** $p < 0.005$.

**Figure S6. sHSP induced during NSPC differentiation promote aggregate formation** *(Related to Figure 6)*

**A,** (***i***) Soluble, misfolded protein substrates (i.e. thermolabile Ubc9$^{ts}$-GFP) are sequestered by yHsp42 into a dynamic protein quality compartment, named Q-bodies. (***ii***) A subset of human sHSPs homologs (indicated in purple) can rescue the misfolded protein sequestration phenotype absent in *hsp42Δ* yeast cells. Epifluorescent images of misfolded Ubc9$^{ts}$-GFP in wild-type and *hsp42Δ* yeast expressing an empty vector (EV), yhsp42, or human sHSPs HSPB1 – HSPB9 as indicated. Graph shows data of cells with inclusions, relative to the WT – empty vector (EV) cells. Quantitative data are presented as mean ± SD of three independent experiments (of 100 cells per condition each). #, significance relative to WT – EV; *, significance relative to *hsp42Δ* – EV (indicated in purple). * $p <0.05$; ** $p < 0.01$; n.s., non-significant.

**B,** Immunoblot analyses of HspB5 in differentiated progeny of control (siCtrl)- and siHspB5-treated NSPCs. HspB5 expression was efficiently depleted by treatment with either a mixed pool of multiple siRNAs (SMARTpool) or single siRNAs (#1 - #4). Gapdh serves as loading control. Degree of HspB5 knockdown is representative for all experiments presented.

**C,** Similar as to differentiated progeny cells of NSPCs treated with the a mixed pool of multiple siRNAs (SMART Pool) targeting HspB5 prior to differentiation, HspB5 cell depletion by various single siRNA-treatment (single siRNA #1 - #4, light gray bars) resulted in a decline in endogenous amyloid-like protein aggregation; quantified by fluorescence (mean ± SD), corrected for the number of cells. Proteostat fluorescence of siCtrl-targeted cells (dark gray bar) is set at 1. Significant inductions in Proteostat fluorescence upon HspB5 depletion are indicated in purple. * $p < 0.05$.

**D,** Confocal imaging of HspB5 (red) and HttQ97-GFP inclusions (green) in differentiated neural cells (single and merged images). Scale bars: 10 µm, trace shows cell outline; arrow indicates HspB5 localization to HttQ97-GFP aggregates.

**E,** (***i***) Confocal imaging of HttQ97-GFP (green) and siGloRed (red; indicator of siRNA transfection) in differentiated progeny cells of control (siCtrl) and siCct1-treated NSPCs. HspB5 expression was depleted by treatment with either a mixed pool of multiple siRNAs (SMART Pool) or single siRNAs (#1 - #4). Scale bars: 10 µm; arrows indicate aggregates, asterisks indicate diffusely expressed HttQ97-GFP. (***ii***) Quantification (mean ± SD) of diffusely localized versus aggregated HttQ97-GFP in differentiated progeny cells of double-transfected siCtrl/siGloRed$^{+}$- and siCct1/siGloRed$^{+}$-treated (SMART Pool *vs.* single siRNAs) NSPCs from at least 40 cells per condition per experiment. Significant inductions in the abundance of HttQ97-GFP inclusions upon HspB5 depletion are indicated in purple ($p < 0.05$).

**Figure S7. Brain aging attenuates TRiC expression, which correlates with an increased deposition of endogenous amyloids, while HspB5 expression remains unaffected** *(Related to Figure 7 and Discussion)*

**A,** Stitched images of endogenous, Proteostat-reactive amyloid deposits (red puncta) in coronal brain sections from young and old mice. DNA stained with Hoechst. Magnification of Proteostat$^{+}$ cells in Dentate Gyrus (*i*), Striatum (*ii*), and Cortex (*iii*) at indicated areas shown in Figure 7B. Scale bars: 500 µm.

**B,** Stitched images of TRiC (Cct1), Sox2 (NPSC marker), and NeuN (neuron marker) in brain sections from young and old mice. Magnification of Cct1 expression at indicated brain areas (i.e. Dentate Gyrus (*i*), Striatum (*ii*,), and Cortex (*iii*)) shown in Figure 7C.

**C, D**, Expression of TRiC subunits (*Cct1-Cct8;* **C**) and sHSPs (*HspB1 – HspB10;* **D**) in NSPCs isolated from young and aged mouse brains (3 and 24 months of age, respectively) and cultured under self-renewal or differentiation (Diff.; differentiated for 7 days) conditions, determined by RT-qPCR. Individual points represent biological replicates (i.e. cells from 3 distinct NSPC isolations), corrected for the expression of the housekeeping genes *Gusb* and *Gapdh*. ****, $p < 0.001$; ***, $p < 0.005$; **, $p < 0.01$; *, $p < 0.05$; n.s., non-significant.

**E,** mRNA expression of *HspB5* in differentiated progeny cells of NSPCs isolated from brains of young adult (3 months old), middle-aged (12 months old), and old (24 months old) mice, analyzed by RT-qPCR. The data represent mean ± SD of three independent experiments with quadruplicate measurements for each condition, corrected for the expression of the housekeeping genes *Gusb* and *Gapdh*. mRNA abundance in differentiated progeny cells of NSPCs isolated from brains of young adult (3 months old) mice is set at 1. n.s., non-significant.

**F,** HspB5 expression in young and aged brain. Confocal imaging of brain sections of the SGZ in the hippocampus of 3 (top) and 24 (bottom) month old mice, immunostained for HspB5 (green) and Sox2 (red; to indicate NSPCs). Scale bars: 100 µm.

### Supplemental Tables

#### Table S1. Overview of disease-linked misfolded proteins studied (*Related to Methods, and Figure 2*)

| Cellular Inclusion | Disease-protein variant | Affected protein | Mutation | Implicated in |
|---|---|---|---|---|
| **Amyloid inclusion** (IPOD/aggresome; *insoluble*) | HttQ97 | Huntingtin | Expansion of polyQ-tract (97Q) in exon 1 | Huntington's Disease (HD) |
| | AR-Q113 | Androgen Receptor | Expansion of polyQ-tract (113Q) | Spinal and Bulbar Muscular Atrophy (SBMA) |
| **Amorphous, non-amyloid aggregate** (JUNQ; *soluble*) | $SOD1^{G37R}$ | Superoxide dismutase 1 | G37R | Amyotrophic Lateral Sclerosis (ALS) |
| | $VHL^{L158P}$ | Von Hippel Lindau | L158P | Cancer |

Table S1. Overview of the disease-linked misfolded proteins studied that have different chaperone requirements and are sequestered into distinct cellular inclusions. IPOD: Insoluble Protein Deposit; JUNQ: Juxtanuclear Quality Control Compartment

#### Table S4. Primers used for RT-qPCR (*Related to Methods; and Figures 7, S4, S7*)

| Gene name | Primer Forward | Primer Reverse |
|---|---|---|
| *HspB1* | CCCGGTTGCCCGATGAGTGG | TCCGCTGACTGCGTGACTGC |
| *HspB2* | CGAGCAGCGCTTCGGAGAAGAT | TGTCCAAATGCCGGCCACCC |
| *HspB3* | CTCCACGGAGAAGCCACCCG | AAGCATTCCCGGGGTGTCCCT |
| *HspB4* | TCGGGCATCTCTGAGGTCCGAT | GGTGAGGCCAGCTCAGGACG |
| *HspB5* | GGATCCGGCGCCCCTTCTTC | TCGTGCTTGCCGTGGACCTC |
| *HspB6* | GCCCAGGCCCAACTTCCGTC | AGAGGTGTCCTGGGTCGGGC |
| *HspB7* | GCCTCAACTCCAGCCGCCAG | ACGAGTGGGTCCAGGCCTCC |
| *HspB8* | GCCAGCGGATGGTTGGGCTC | GCGGGAGGAGAGCGGTGAGT |
| *HspB9* | CCAGCGTGGCCCTTGCTGAA | GGTCGGCGGGAGCTGCATTT |
| *HspB10* | ACTCCTACGGGCTCGGCAGC | GGGCTCTGGGAAGGGGACGAG |
| *Cct1* | GGGTCCTTTGTCCGTGTTC | CTGGCCCAAAAGAACTTTTAAC |
| *Cct2* | CCCGCAGCAAAGGTTCTA | TTGCAATTAAAGATTCTGCTTCC |
| *Cct3* | GGACCCAGGATGAAGAGGTT | TGCATCTGCTGCTCTAGGAA |
| *Cct4* | TCTGTGTCATCCGATGCTTAGT | GGCCAGCTCTATTTCTGGAG |
| *Cct5* | CTGTTTGCACAAGGGCAGTA | CCATCTGGGTGGCAAGAG |
| *Cct6* | GTGTGGGATAACTACTGTGTGAAG A | CAGCTCGCATGATCTCGTC |
| *Cct7* | GCTGTGACTGTGAAGAAGCAA | ACGGCATCAACCACCATC |
| *Cct8* | GGCCTCTCATATGCAAGAACA | GCGAACACCAGAACGAAGTT |
| *Gusb* | GATGTGGTCTGTGGCCAAT | TGTGGGTGATCAGCGTCTT |
| *Gapdh* | GGAGCGAGACCCCACTAACA | ACATACTCAGCACCGGCCTC |

Table S4. Primers used for RT-qPCR

**Data S1. Uncropped Images of Immunoblots** *(Related to Figures 2, 3, 5, and 6)*

**A.** Immunoblot for Ubiquitin in NSPCs at basal or proteasome-inhibited conditions (MG132; 15 µM, 4 h prior to fixation). Gapdh serves as loading control. Red boxes indicate lanes shown in Figure 2H*ii.* (*Related to Figure 2Hii)*

**B.** Immunoblots for indicated chaperones for three independent samples of NSPCs and early, intermediate, and fully differentiated cells. Gapdh serves as loading control. Red boxes indicate blot area shown in Figure 3E*i. (Related to Figure 3E)*

**C, D.** Immunoblots for all 8 Cct subunits of the TRiC chaperonin complex (**C**) or various sHsps (**D**) for three independent samples of NSPCs and progeny cells. Gapdh (**C**) and actin (**D**) serve as loading controls. Red boxes indicate blot area shown in Figures *(C, Related to Figure 4A; D, Related to Figure 4B)*

**E.** Immunoblot analyses of TRiC (Cct1), NSPC markers (Nestin, Sox2), and neuronal marker (NeuN). Gapdh serves as loading control. Red boxes indicate blot area shown in Figure. *(Related to Figure 5A)*

**F.** Coomassie-stained SDS-PAGE gel and Cct1 immunoblot in siCtrl and siCct1-treated NSPCs. Gapdh serves as loading control. Red boxes indicate lanes/area shown in Figure. *(Related to Figure 5D)*

**G.** Immunoblot analysis with antibodies for Psma1-7 (20S catalytic core subunit), Psmd2 (19S regulatory subunit), and Gapdh (loading control) in NSPCs and differentiated progeny. Red boxes indicate blot area shown in Figure. *(Related to Figures 2; S2G)*

**H.** Immunoblot for LC3 and Gapdh as indicated. Red boxes indicate blot area shown in Figure. *(Related to Figures 3; S3G)*

**I.** Immunoblot analyses of Cct1 and Gapdh in control (siCtrl)- and siCct1-targeted NSPCs. Red boxes indicate blot area shown in Figure. *(Related to Figures 5; S5A)*

**J.** Immunoblot analyses of Cct2 and Gapdh in control (siCtrl)- and siCct2-targeted NSPCs. Red boxes indicate blot area shown in Figure. *(Related to Figures 5; S5E)*

**K.** Immunoblot analyses of HspB5 and Gapdh in differentiated progeny of control (siCtrl)- and siHspB5-treated NSPCs. Red boxes indicate blot area shown in Figure. *(Related to Figures 6; S6B)*

**Data S1. Uncropped images of Immunoblots**

**A.**

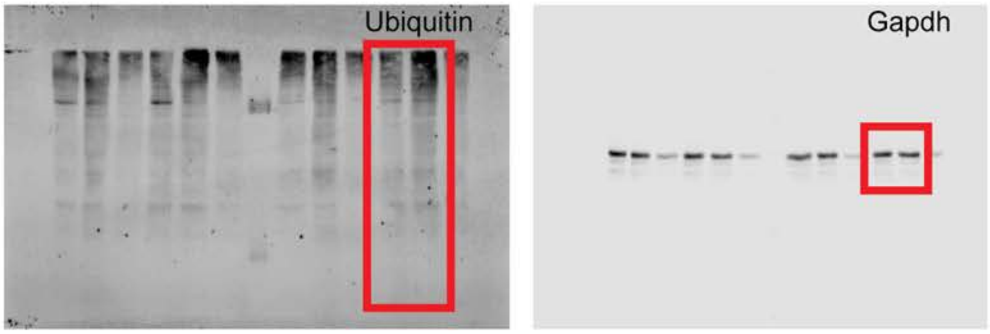


**B.**

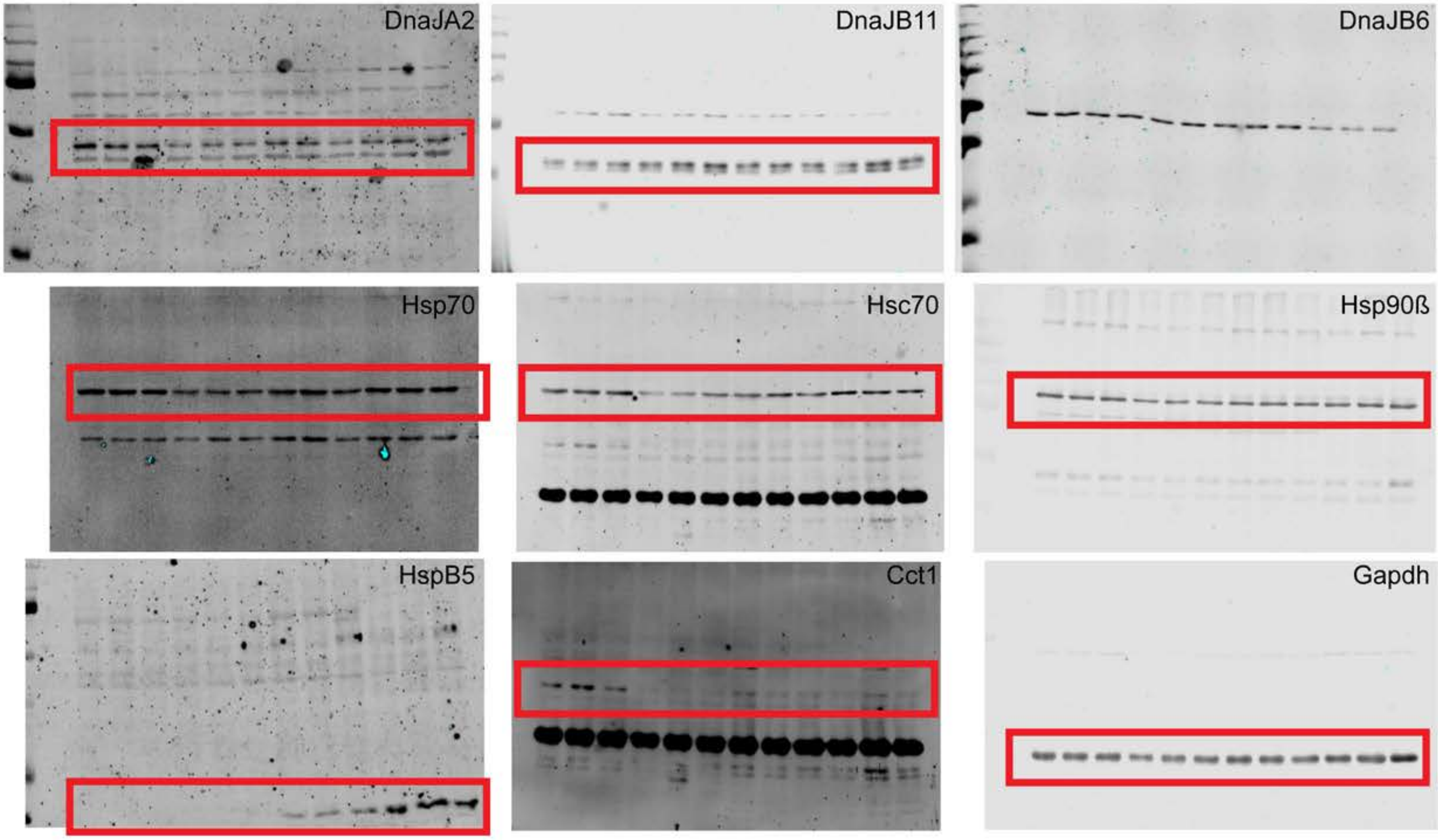

C.

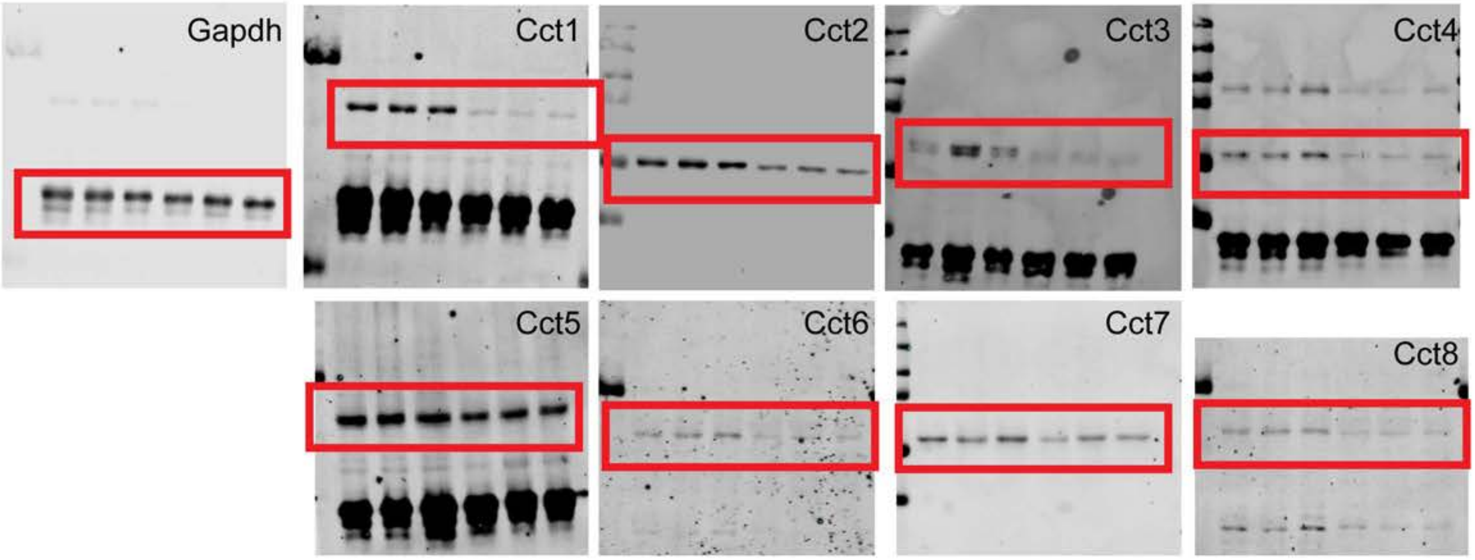
Gapdh
Cct1
Cct2
Cct3
Cct4
Cct5
Cct6
Cct7
Cct8

D.

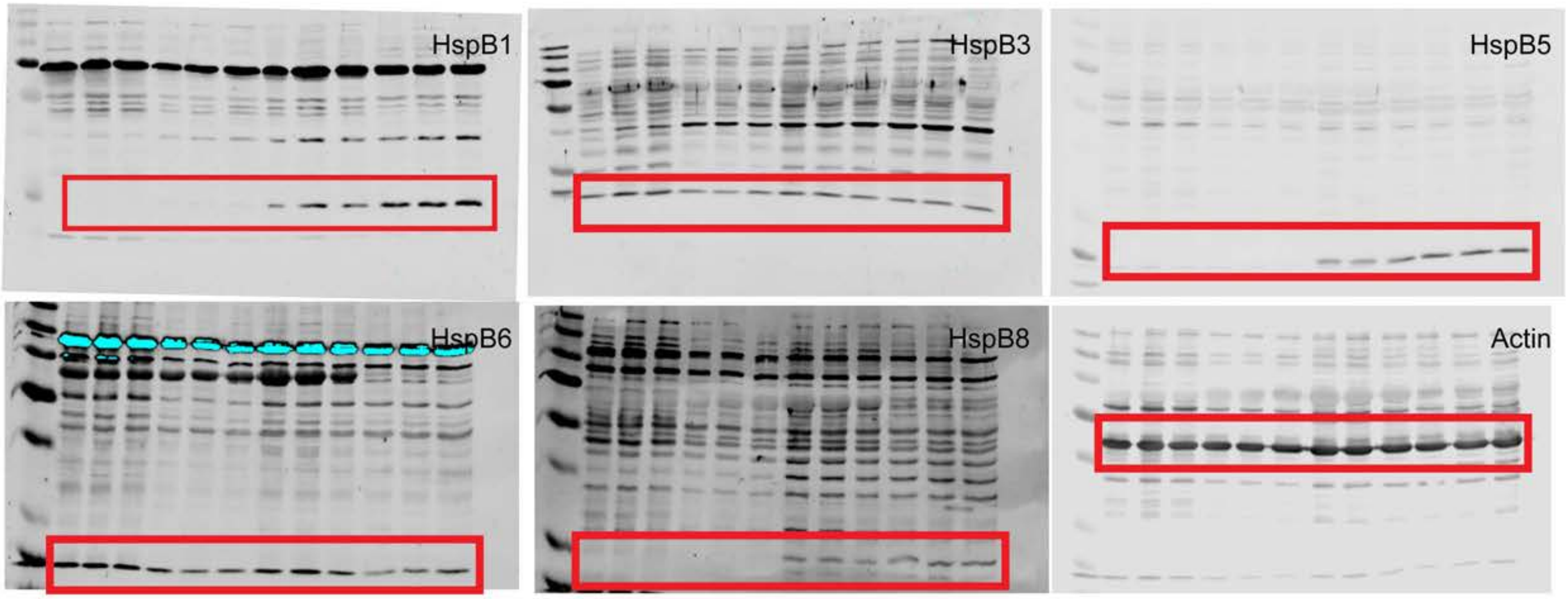
HspB1
HspB3
HspB5
HspB6
HspB8
Actin

E.

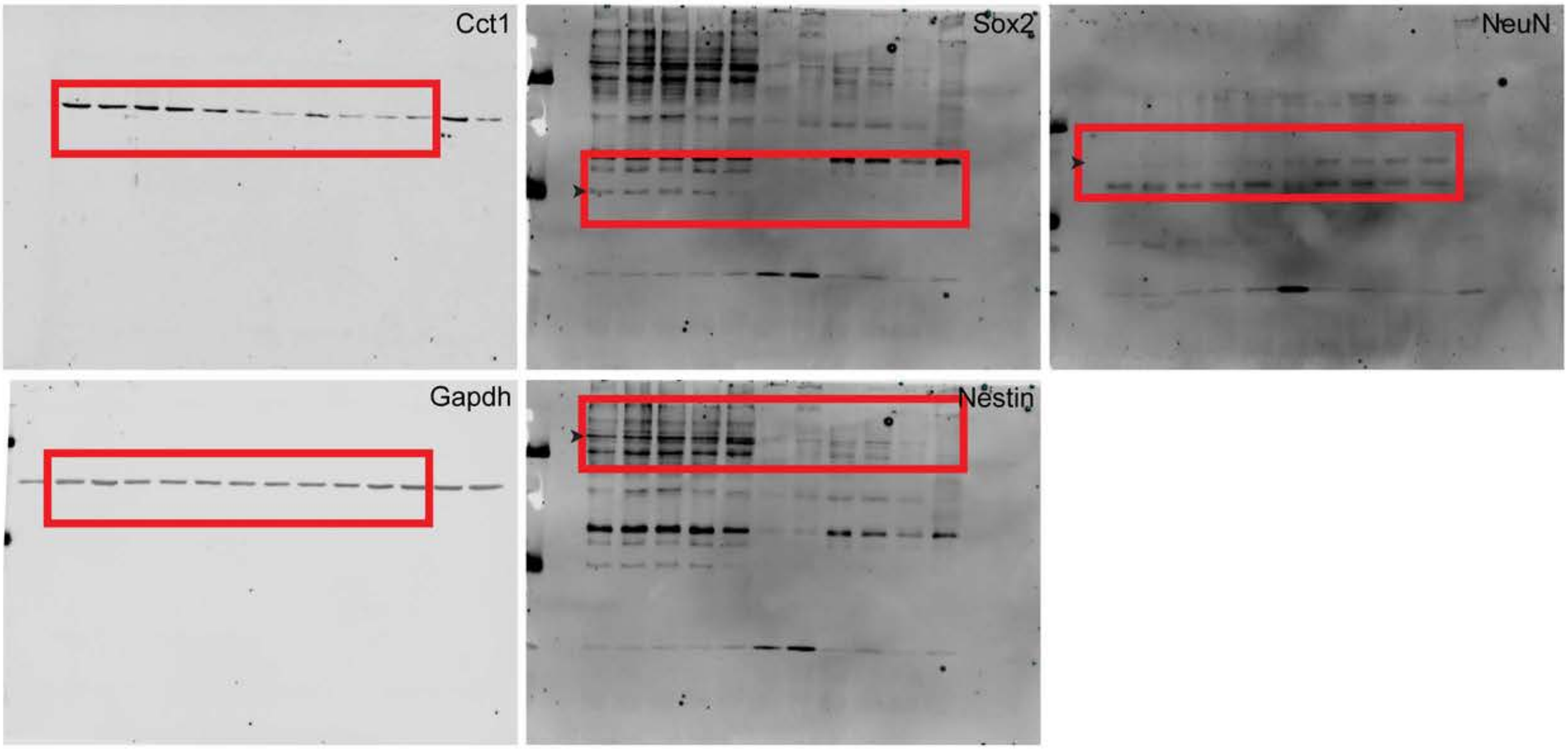


F.

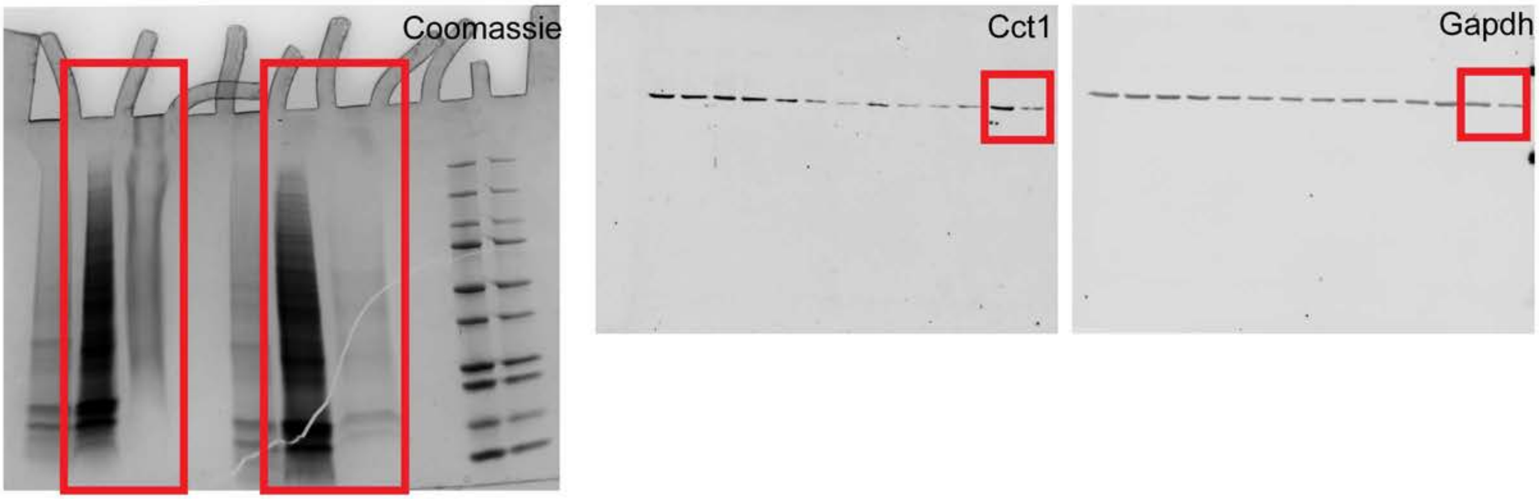

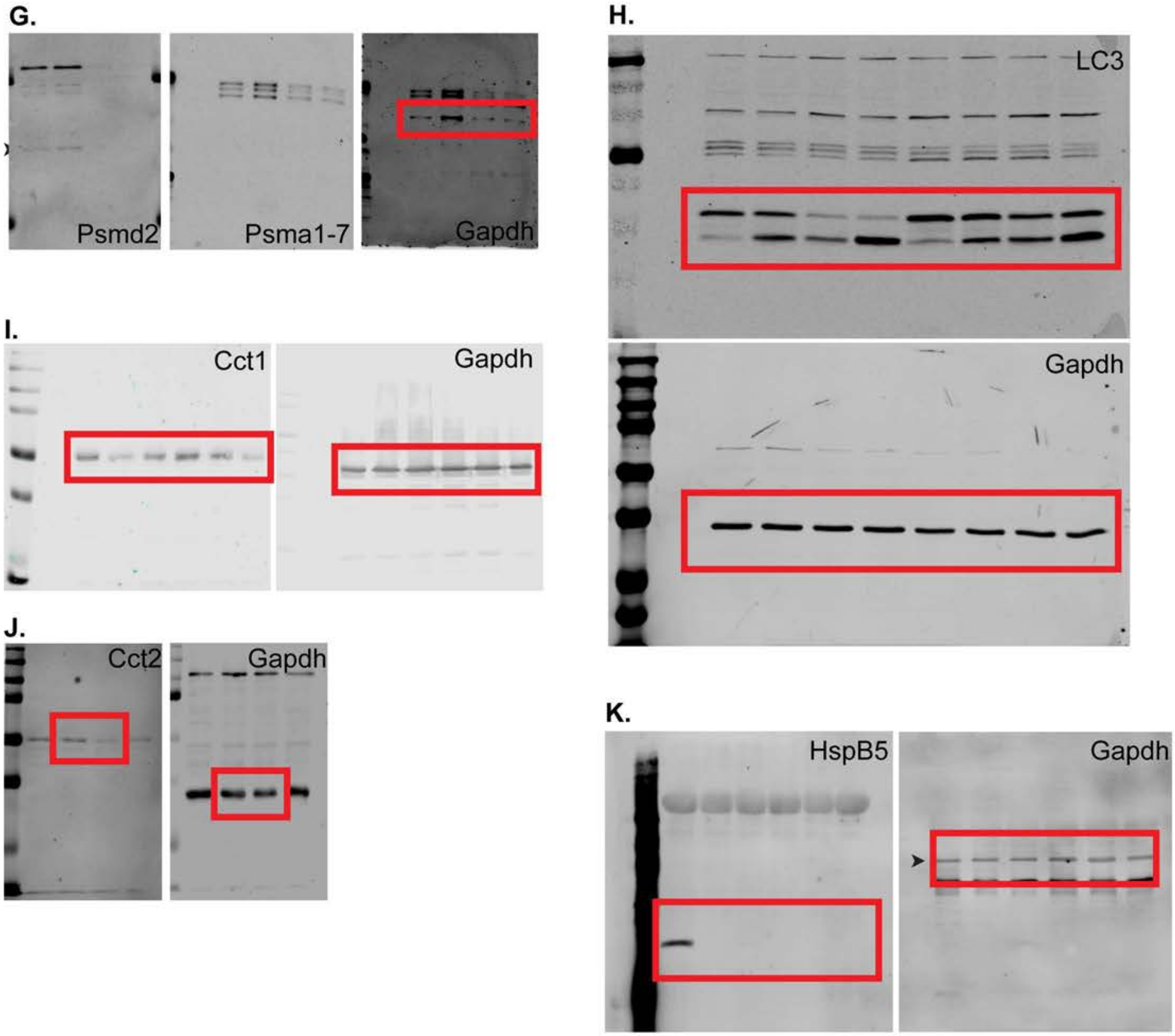

G.
Psmd2
Psma1-7
Gapdh
H.
LC3
Gapdh
I.
Cct1
Gapdh
J.
Cct2
Gapdh
K.
HspB5
Gapdh
